# PI3K block restores age-dependent neurovascular coupling defects associated with cerebral small vessel disease

**DOI:** 10.1101/2023.03.03.531032

**Authors:** Pratish Thakore, Evan Yamasaki, Sher Ali, Alfredo Sanchez Solano, Cassandre Labelle-Dumais, Xiao Gao, Myriam M. Chaumeil, Douglas B. Gould, Scott Earley

## Abstract

Neurovascular coupling (NVC), a vital physiological process that rapidly and precisely directs localized blood flow to the most active regions of the brain, is accomplished in part by the vast network of cerebral capillaries acting as a sensory web capable of detecting increases in neuronal activity and orchestrating the dilation of upstream parenchymal arterioles. Here, we report a *Col4a1* mutant mouse model of cerebral small vessel disease (cSVD) with age-dependent defects in capillary-to-arteriole dilation, functional hyperemia in the brain, and memory. The fundamental defect in aged mutant animals was the depletion of the minor membrane phospholipid phosphatidylinositol 4,5 bisphosphate (PIP_2_) in brain capillary endothelial cells, leading to the loss of inwardly rectifier K^+^ (Kir2.1) channel activity. Blocking phosphatidylinositol-3-kinase (PI3K), an enzyme that diminishes the bioavailability of PIP_2_ by converting it to phosphatidylinositol (3,4,5)-trisphosphate (PIP_3_), restored Kir2.1 channel activity, capillary-to-arteriole dilation, and functional hyperemia. In longitudinal studies, chronic PI3K inhibition also improved the memory function of aged *Col4a1* mutant mice. Our data suggest that PI3K inhibition is a viable therapeutic strategy for treating defective NVC and cognitive impairment associated with cSVD.

**One-sentence summary:** PI3K inhibition rescues neurovascular coupling defects in cerebral small vessel disease.

## Introduction

A group of familial and idiopathic pathologies known as cerebral small vessel diseases (cSVDs) are a leading cause of vascular contributions to cognitive impairment and dementia (VCID) ^1^, second only to Alzheimer’s disease as the most common cause of cognitive impairment in adults ^2,3^. The global impact of cSVDs is massive and rapidly growing as the world’s population ages ^4^, but little is currently known about the pathogenesis of the disease and no specific treatments exist. Brain atrophy, lacunes, enlarged perivascular spaces, intracerebral hemorrhages (ICH), white matter hyperintensities, and microinfarcts detected by magnetic resonance imaging (MRI) are clinical signs of irreversible brain damage brought on by cSVDs ^5^. cSVDs also impair functional hyperemia, a crucial physiological process in which local blood flow is swiftly and precisely diverted to the most active brain regions ^6-9^. Dysregulation of cerebral blood flow contributes to vascular dementia ^1^, but it is not known if resolution of this impairment can delay or reverse cognitive decline. Autosomal dominant mutations in the genes encoding collagen type IV alpha 1 (COL4A1) and alpha 2 (COL4A2) cause an inherited form of cSVD as part of a multisystem disorder called Gould syndrome ^10-13^. In this study, we used *Col4a1* mutant mice to elucidate the molecular events that disrupt functional hyperemia in this type of cSVD and applied these findings to repair the deficit.

Functional hyperemia is accomplished by a collection of physiological processes called neurovascular coupling (NVC). During NVC, active neurons trigger the dilation of upstream pial arteries and parenchymal arterioles supplying the brain, boosting blood flow to fulfill regional metabolic demands ^14^. The underlying mechanisms have been intensely studied for decades ^15-17^, however much remains to be discovered. A new paradigm has emerged envisioning the vast cerebral capillary network as a sensory web capable of detecting increases in neuronal activity and orchestrating the dilation of upstream parenchymal arterioles ^18^. In this arrangement, substances released from nearby active neurons and/or astrocytic endfeet stimulate receptors on brain capillary endothelial cells (ECs) to generate retrograde propagating vasodilator signals that travel against the flow of blood and dilate upstream arterioles. Two types of ion channels present on brain capillary ECs, inward-rectifying K^+^ (Kir2.1) channels, and transient receptor potential ankyrin 1 (TRPA1) cation channels, are critical sensors of neuronal activity and are necessary for NVC and functional hyperemia in the brain ^18,19^. Kir2.1 channels are activated by K^+^ ions released during neuronal activity and initiate rapid retrograde propagating electrical signals that ultimately dilate upstream arterioles and increase blood flow to meet neuronal demand ^18,20,21^. TRPA1 channels are activated by reactive oxygen species metabolites and stimulate propagating intercellular Ca^2+^ waves that dilate upstream arterioles ^19^. Capillary-to-arteriole dilation is impaired in the CADASIL (Cerebral Autosomal Dominant Arteriopathy with Sub-cortical Infarcts and Leukoencephalopathy) cSVD mouse model ^22^, but the impact of *Col4a1* mutations on this process, and NVC coupling, is unknown.

Here, we report that Kir2.1 channel activity in ECs of cerebral arteries and brain capillaries from *Col4a1* mutant mice is lost during aging. K^+^-induced capillary-to-arteriole dilation was absent in vascular preparations from 12-month (M)-old mutant animals, resulting in impaired functional hyperemia in the somatosensory cortex and memory deficits. Reduced Kir2.1 channel activity was due to diminished phosphatidylinositol 4,5 bisphosphate (PIP_2_), a minor membrane phospholipid that is an essential co-factor for Kir2.1 activity ^23,24^. Kir2.1 channel activity and capillary-to-arteriole dilation was restored by blocking phosphoinositide 3-kinase (PI3K), an enzyme that converts PIP_2_ to phosphatidylinositol (3,4,5)-trisphosphate (PIP_3_). Chronic treatment of mutant mice with a PI3K antagonist restored Kir2.1 channel activity and capillary-to-arteriole dilation, improved functional hyperemia, and largely resolved memory deficits. Our findings reveal a novel age-dependent mechanism of impaired NVC associated with cSVD and identify PI3K as a potential therapeutic target for treating vascular cognitive impairment and dementia.

## Results

### Age-dependent loss of Kir2.1 channel activity and capillary-to-arteriole dilation in Col4a1^+/G394V^ mice

Mutant mice used for this study harbor a point mutation in one copy of the *Col4a1* gene that results in the glycine (G) to valine (V) substitution at position 394 of the COL4A1 polypeptide (*Col4a1^+/G394V^*) ^25^. Wild type littermates were used as controls for all experiments. Both male and female mice were studied, and no sex-specific differences were observed. Age is the most critical risk factor for cSVD and vascular dementia ^26,27^. Therefore, we used *Col4a1^+/G394V^* and control mice at 3 and 12 months (M) of age, representing young adulthood and middle age, respectively, throughout this study.

Using whole-cell patch clamp electrophysiology, we first investigated how Kir2.1 currents in freshly isolated brain capillary ECs (Fig. 1A) were affected by *Col4a1^G394V^* mutation. Kir2.1 channels were activated by increasing the [K^+^] of the bath solution to 60 mM, and currents were recorded as voltage ramps (−100 to +40 mV) were applied. The selective Kir channel blocker BaCl_2_ (10 μM) was used to isolate the current. Kir2.1 current densities in brain capillary ECs from young adult mice did not differ between mutant and control mice (Fig. 1B) but were significantly reduced in brain capillary ECs from 12 M-old *Col4a1^+^*^/^*^G394V^* mice compared with aged-matched control animals (Fig. 1C). We then utilized an innovative *ex vivo* cerebral microvascular preparation in which parenchymal arteriole segments with intact capillary branches were isolated from the brain, cannulated and pressurized ^18,19^ to determine if loss of Kir2.1 channel activity translated into impaired capillary-to-arteriole dilation. Capillaries were stimulated by locally applying pulses (7 seconds; 10 psi) of KCl (10 mM) dissolved in artificial cerebral spinal fluid (aCSF) using a micropipette attached to a picospritzer (Fig. 1D). Focal application of K^+^ to the capillary beds of 3 M-old *Col4a1^+/+^* and *Col4a1^+/G394V^* mice produced robust, reversible, and reproducible dilations of the upstream arterioles (Fig. 1E and F). The addition of the nitric oxide donor sodium nitroprusside (SNP) to the tissue bath maximally dilated arterioles from both groups (Fig. 1G). Similar responses were observed when tissues from 12 M-old control mice were used (Fig. 1H and I). In contrast, focal application of K^+^ onto capillaries from 12 M-old *Col4a1^+/G394V^* mice did not dilate upstream arterioles. However, SNP produced maximal dilation, demonstrating the viability of the preparation (Fig. 1J). These results show that the age-dependent impairment of Kir2.1 current activity in brain capillary ECs from *Col4a1^+/G394V^* mice eliminates capillary-to-arteriole dilation.

**Fig. 1.**
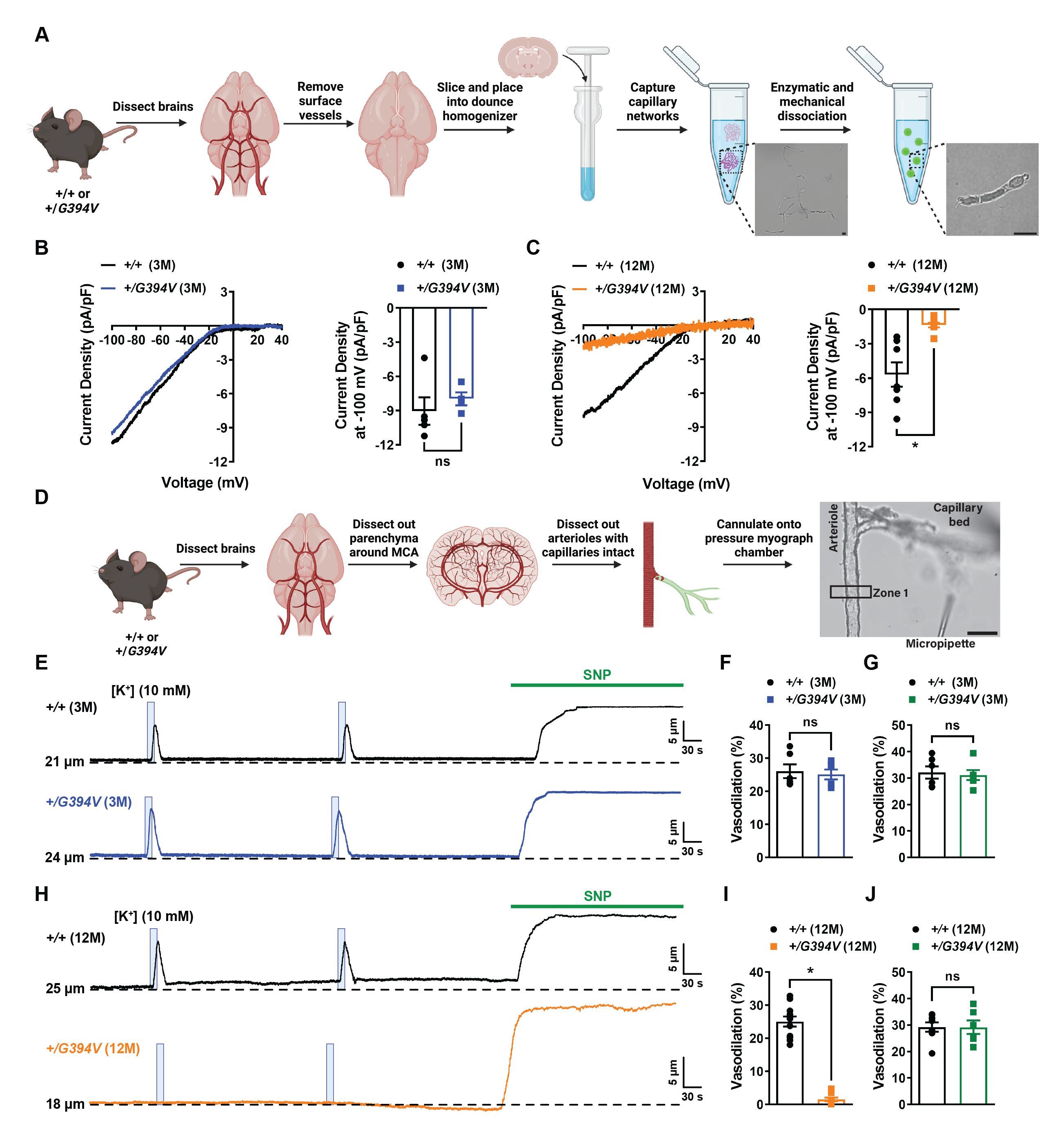
Age-dependent loss of Kir2.1 channel activity and capillary-to-arteriole dilation in Col4a1^+/G394V^ mice. (A) Illustration of the brain capillary EC isolation procedure. Scale bar = 10 µm. (B) Representative I-V traces and summary data showing Kir2.1 current densities in freshly isolated capillary ECs from 3 M-old *Col4a1^+/+^* and *Col4a1^+/G394V^* mice (n = 4–5 cells from 4 to 5 animals per group, ns = not significant, unpaired t-test). (C) Representative I-V traces and summary data showing Kir2.1 current densities in freshly isolated capillary ECs from 12 M-old *Col4a1^+/+^* and *Col4a1^+/G394V^* mice (n = 7–8 cells from 4 animals per group; *p<0.05, unpaired t-test). (D) Illustration of the microvascular preparation. Parenchymal arterioles with intact capillaries were carefully dissected and cannulated onto a pressure myograph chamber, and compounds of interest were focally applied to capillary extremities. Scale bar = 50 µm. (E and F) Representative traces (E) and summary data (F) showing K^+^ (10 mM, blue box)-induced dilation of upstream arterioles in preparations from 3 M-old *Col4a1^+/+^* and *Col4a1^+/G394V^* mice (n = 6 preparations from 3 animals per group, ns = not significant, unpaired t-test). (G) The dilation produced by superfusing SNP (10 µM) in preparations from 3 M-old *Col4a1^+/+^* and *Col4a1^+/G394V^* mice (n = 6 preparations from 3 animals per group, ns = not significant, unpaired t-test). (H and I) Representative traces (H) and summary data (I) showing K^+^ (10 mM, blue box)-induced dilation of upstream arterioles in preparations from 12 M-old *Col4a1^+/+^* and *Col4a1^+/G394V^* mice (n = 11 preparations from 6 to 7 animals per group, *p<0.05, unpaired t-test). (J) The dilation produced by superfusing SNP (10 µM) in preparations from 12 M-old *Col4a1^+/+^* and *Col4a1^+/G394V^* mice (n = 6–8 preparations from 3 to 5 animals per group, ns = not significant, unpaired t-test).

We also investigated the effects of the *Col4a1* mutation on Kir2.1 channel activity in the ECs that line cerebral arteries (arteriolar ECs) (Fig. 1-supplement 1A) and found that these currents did not differ for 3 M-old mice but were blunted in 12 M-old *Col4a1^+/G394V^* animals compared to age-matched controls (Fig. 1-supplement 1B and C). Prior studies show that cerebral arteries dilate when [K^+^] is raised from 3 mM to between 8 and 20 mM ^28,29^ because increased Kir2.1 channel activity hyperpolarizes the membrane potential of vascular smooth muscle. Higher [K^+^] (greater than 30 mM) collapses the gradient, causing membrane depolarization and vasoconstriction. To determine the functional consequence of reduced Kir2.1 channel activity in arteriolar ECs from *Col4a1* mutants, we used standard pressure myography techniques ^30^ to investigate the effects of increasing [K^+^] on vasomotor responses of isolated cerebral arteries. We found that raising the external [K^+^] from 3 mM to a range of 8 to 20 mM dilated cerebral arteries from 12-M old control animals but had no effect on arteries from 12 M-old *Col4a1^+^*^/^*^G394V^* mice (Fig. 1 – supplement 1D and E). In contrast, vasoconstriction in response to higher [K^+^] (30 and 60 mM) did not differ between 12 M-old *Col4a1^+^*^/^*^G394V^* and control mice (Fig. 1 – supplement 1F). These results demonstrate that age-dependent loss of Kir2.1 currents impairs K^+^-induced vasodilation. In addition, our data show that fundamental voltage-dependent contractile mechanisms are not grossly affected.

Prior studies have demonstrated mild ICHs in the brains of 12 M-old *Col4a1^+/G394V^* mice ^31,32^. We further demonstrated this using *in vivo* susceptibility-weighted magnetic resonance imaging (SWI) to identify possible brain lesions. No hypointense lesions were detected on any SWI images from 12 M-old *Col4a1^+/+^* and *Col4a1^+^*^/^*^G394V^* mice (Fig. 1 – supplement 2A), demonstrating that this mutation does not induce magnetic resonance (MR)-detectable hemorrhagic lesions. Volumetric analysis of the T2-weighted (T2W) images showed that the ventricle/brain ratio was not significantly different between 12 M-old *Col4a1^+/+^* and *Col4a1^+/G394V^* mice (Fig. 1 – supplement 2B), further demonstrating that these mutant mice do not show obvious pathology in these imaging modalities.

### Age-dependent impairment of functional hyperemia in Col4a1^+/G394V^ mice

The effects of the *Col4a1^G394V^* mutation on brain hemodynamics *in vivo* were investigated using the thinned-skull laser Doppler flowmetry method to measure blood flow changes in the somatosensory cortex in response to stimulating whiskers for 1, 2, or 5 seconds (s) (Fig. 2A and B) ^19,33,34^. Stimulating contralateral whiskers for 1 s reproducibly increased blood flow in the somatosensory cortex of 3 M-old mice, with no differences in the amplitude, latency, or kinetics of the response between control and mutant mice (Fig. 2C to H). The magnitude of blood flow increases in response to 1 s stimulation were significantly blunted in 12 M-old *Col4a1^+/G394V^* mice compared to age-matched controls (Fig. 2I and J). In addition, the latency of the blood flow response was increased, and the rise and decay rates were diminished in 12 M-old mutant mice (Fig. 2K to N). Similar outcomes were observed for the 2 s (Fig. 2 – supplement 1), and 5 s (Fig. 2O to Z) stimulation protocols – no differences were detected in 3 M-old mice, but a blunted increases in blood flow, increased latency, and decreased rise rate and decay rate was detected for 12 M-old *Col4a1^+/G394V^* mice compared with controls. In addition, the duration of the blood flow increase was reduced in 12-M old mutant mice compared with controls in the 5 s stimulation protocol. Stimulation of ipsilateral whiskers failed to produce any change in the blood flow (Fig. 2 – supplement 2). These results demonstrate that loss of brain EC Kir2.1 channel activity and capillary-to-arteriole dilation is associated with impaired functional hyperemia in 12 M-old *Col4a1^+/G394V^* mice *in vivo*.

**Fig. 2.**
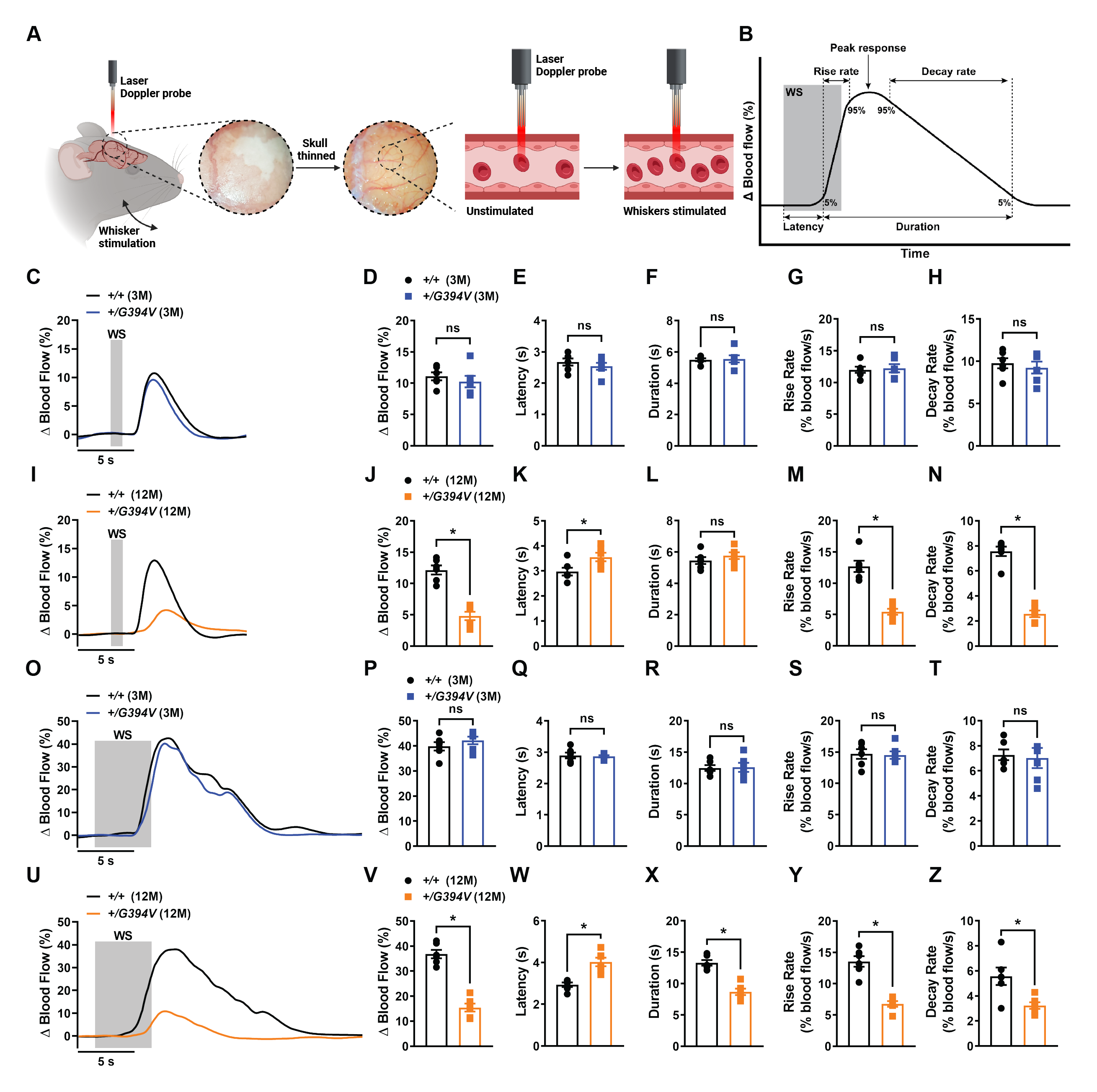
Age-dependent impairment of functional hyperemia in Col4a1^+/G394V^ mice. (A) Illustration demonstrating the functional hyperemia assessment procedure in the mouse somatosensory cortex. (B) Illustration demonstrating the parameters that were analyzed. (C and D) Representative traces (C) and summary data (D) showing the increase in blood flow following 1 s contralateral whisker stimulation (WS) in 3 M-old *Col4a1^+/+^* and *Col4a1^+/G394V^* mice (n = 6 animals per group, ns = not significant, unpaired t-test). (E to H) Latency (E), duration (F), rise rate (G), and decay rate (H) were also analyzed (n= 6 animals per group, ns = not significant, unpaired t-test). (I and J) Representative traces (I) and summary data (J) showing the increase in blood flow following 1 s contralateral WS in 12 M-old *Col4a1^+/+^* and *Col4a1^+/G394V^* mice (n = 6 animals per group, *p<0.05, unpaired t-test). (K to N) Latency (K), duration (L), rise rate (M), and decay rate (N) were also analyzed (n = 6 animals per group, *p<0.05, ns = not significant, unpaired t-test). (O and P) Representative traces (O) and summary data (P) showing the increase in blood flow following 5 s contralateral WS in 3 M-old *Col4a1^+/+^* and *Col4a1^+/G394V^* mice (n = 6 animals per group, ns = not significant, unpaired t-test). (Q to T) Latency (Q), duration (R), rise rate (S), and decay rate (T) were also analyzed (n= 6 animals per group, ns = not significant, unpaired t-test). (U and V) Representative traces (U) and summary data (V) showing the increase in blood flow following 5 s contralateral WS in 12 M-old *Col4a1^+/+^* and *Col4a1^+/G394V^* mice (n = 6 animals per group, *p<0.05, unpaired t-test). (W to Z) Latency (W), duration (X), rise rate (Y), and decay rate (Z) were also analyzed (n = 6 animals per group, *p<0.05, unpaired t-test).

### Age-dependent memory deficits in Col4a1^+/G394V^ mice

Impaired NVC is associated with deficiencies in spatial working and recognition memory ^35,36^. Here, the spontaneous alternation behavioral assay was used to determine if *Col4a1^+/G394V^* mice develop memory deficits. Mice were freely allowed to explore all three arms of a Y-shaped maze for 10 minutes, and their consecutive entries into each of the arms were recorded and reported as % alternation (Fig. 3A) ^37,38^. Spontaneous alternation, the maximum number of alternations, and the total distance moved did not differ between 3 M-old *Col4a1^+/G394V^* and control mice (Fig. 3B to D). However, spontaneous alternation was significantly diminished for 12 M-old *Col4a1^+/G394V^* mice compared with age-matched controls (Fig. 3E), suggesting deficits in spatial working memory. The maximum alternation and total distance moved did not differ between 12 M-old mutant and control mice, indicating that impaired mobility does not account for reduced alternation (Fig. 3F and G).

**Fig. 3.**
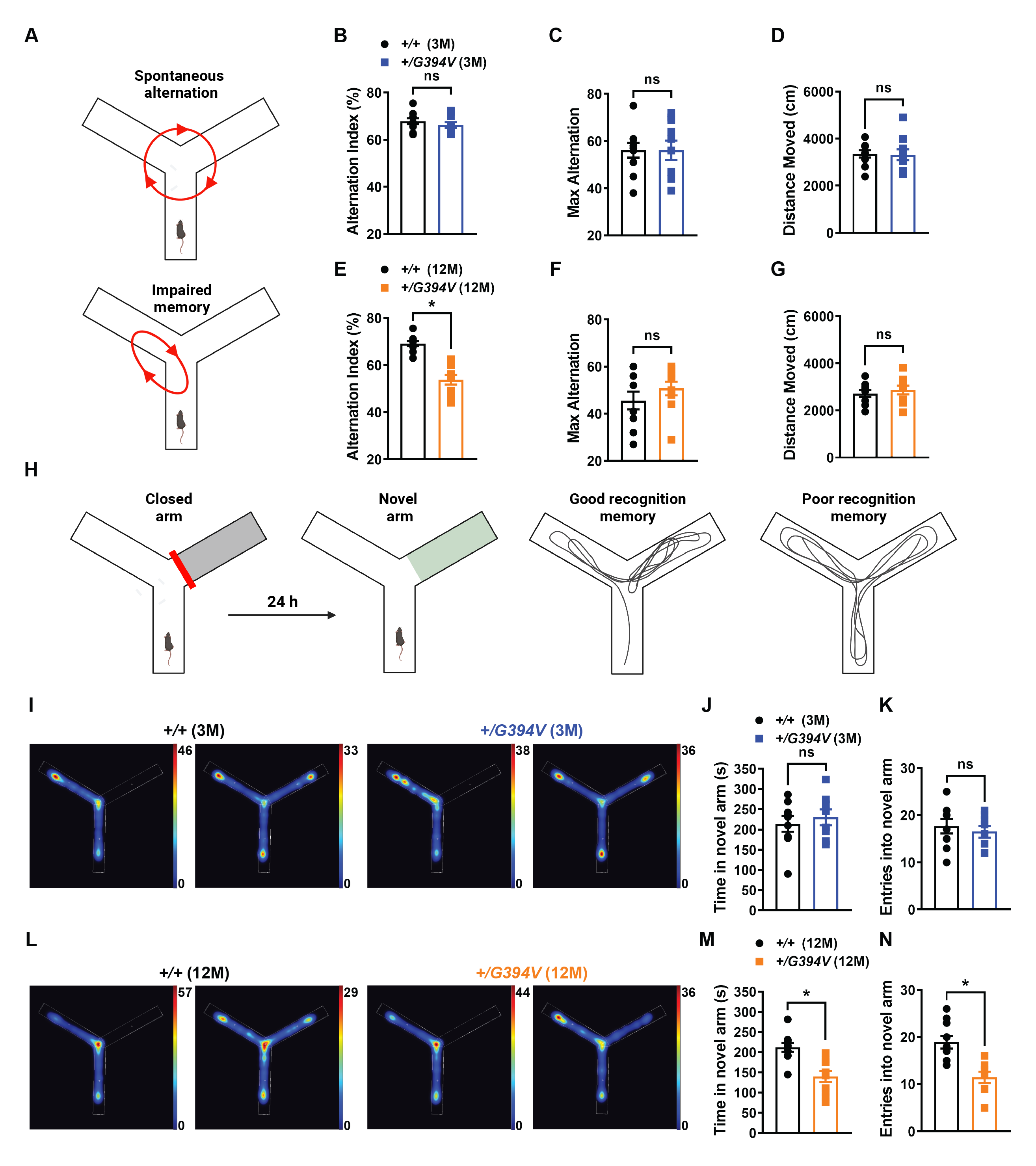
Age-dependent memory deficits in Col4a1^+/G394V^ mice. (A) Illustration of the Y-maze spontaneous alternation behavior assay showing examples of a spontaneous (top) and nonspontaneous (bottom) alternation. (B) Summary data showing alternation index, an indicator spatial working memory, in 3 M-old *Col4a1*^+/+^ and *Col4a1*^+/G394V^ mice (n = 10 animals per group, ns = not significant, unpaired t-test). (C and D) Summary data showing max alternation (C) and distance moved (D), indicative of exploratory activity, in 3 M-old *Col4a1*^+/+^ and *Col4a1*^+/G394V^ mice (n = 10 animals per group, ns = not significant, unpaired t-test). (E) Summary data showing the alternation index of 12 M-old *Col4a1*^+/+^ and *Col4a1*^+/G394V^ mice (n = 10 animals per group, *p<0.05, unpaired t-test). (F and G) Summary data showing max alternation (F) and distance moved (G) in 12 M-old *Col4a1*^+/+^ and *Col4a1*^+/G394V^ mice (n = 10 animals per group, ns = not significant, unpaired t-test). (H) Illustration demonstrating typical and impaired Y-maze novel arm behavior. (I) Representative heatmaps showing the time (s) 3 M-old *Col4a1*^+/+^ and *Col4a1*^+/G394V^ mice spent in areas of the Y-maze during the novel arm test. (J and K) Summary data showing the time spent (J) and entries (K) into the novel arm (n = 8–9 animals per group, ns = not significant, unpaired t-test). (L) Representative heatmaps showing the time (s) 12 M-old *Col4a1*^+/+^ and *Col4a1*^+/G394V^ mice spent in areas of the Y-maze during the novel arm test. (M and N) Summary data showing the time spent (M) and entries (N) into the novel arm (n = 10 animals per group, *p<0.05, unpaired t-test).

A second behavioral assay, the novel arm test, was also used to evaluate recognition memory function in mutant mice. This task is driven by the innate curiosity of mice to explore previously unvisited areas ^39,40^. Initially, mice were allowed to explore a Y-maze for 10 minutes with one arm of the maze blocked. After 24 hours, the mice were allowed to explore the maze for 10 minutes with all arms open (Fig. 3H). Mice with normal recognition memory spend more time exploring the novel arm than those explored on the previous day. The dwell time and the number of entries in the novel arm did not differ between 3 M-old *Col4a1^+/G394V^* and control mice (Fig. 3I to K). In contrast, 12 M-old *Col4a1^+/G394V^* mice spent significantly less time and had fewer entries into the novel arm compared with age-matched controls (Fig. 3L to N). These data provide further evidence of age-dependent memory deficits in *Col4a1^+/G394V^* mice.

### PIP_2_ depletion reduces Kir2.1 currents in 12 M-old Col4a1^+/G394V^ mice

Kir2.1 channels require PIP_2_ for activity ^23,24^, and prior studies reported that pathogenic loss of PIP_2_ diminished Kir2.1 channel activity in brain capillary ECs from CADASIL cSVD mice and 5xFAD familial Alzheimer’s disease mice ^22,41^. Therefore, we investigated the possibility that PIP_2_ depletion also reduces Kir2.1 current density in ECs from 12 M-old *Col4a1^+/G394V^* mice. When exogenous diC8-PIP_2_ (10 µM) was included in the intracellular solution during whole-cell patch-clamp experiments, Kir2.1 current density in arteriolar ECs (Fig. 4A) and brain capillary EC (Fig. 4B) did not differ between 12 M-old control and age-matched *Col4a1^+/G394V^* mice, suggesting that PIP_2_ depletion is responsible for Kir2.1 current deficiencies in the mutant mice. The steady-state amount of PIP_2_ present in the inner leaflet of the plasma membrane is determined by the relative rates of PIP_2_ synthesis and degradation. PIP_2_ is synthesized by the sequential action of phosphatidylinositol 4-kinase (PI4K), an enzyme that converts phosphatidylinositol (PI) to phosphatidylinositol 4-phosphate (PIP), and phosphatidylinositol 4-phosphate 5-kinase (PIP5K), which converts PIP to PIP_2_. PI4K and PIP5K activity require high levels of ATP (i.e., the K_M_ of PI4K for ATP is ∼0.4 to 1 mM) (Fig. 4 – supplement 1A) ^42-44^. PIP_2_ deficiency in capillary ECs from CADASIL mice was attributed to reduced ATP levels and diminished synthesis ^22^. We measured ATP in isolated brain capillaries using a luciferase-based assay and found that ATP levels did not differ between 12 M-old *Col4a1^+/G394V^* and control mice (Fig. 4 – supplement 1B), suggesting that PIP_2_ synthesis is not impaired by lack of ATP in *Col4a1* mutants. Hydrolysis of PIP_2_ by phospholipase C (PLC) to form inositol trisphosphate (IP_3_) and diacylglycerol (DAG) diminishes steady-state PIP_2_ levels (Fig. 4 – supplement 1C) ^45^. However, blocking PLC activity with U73122 (10 µM) did not restore Kir2.1 currents in capillary ECs from 12 M-old *Col4a1^+/G394V^* mice, suggesting that PIP_2_ depletion does not result from elevated levels of PLC activity (Fig. 4 – supplement 1D). PIP_2_ bioavailability is reduced when the enzyme PI3K phosphorylates it to PIP_3_ (Fig. 4C). When PI3K was blocked using the selective inhibitor GSK1059615 (10 nM), Kir2.1 currents in capillary ECs from 12 M-old *Col4a1^+/G394V^* mice were restored to control levels (Fig. 4D), suggesting that PIP_2_ depletion in mutant mice results from elevated PI3K activity.

**Fig. 4.**
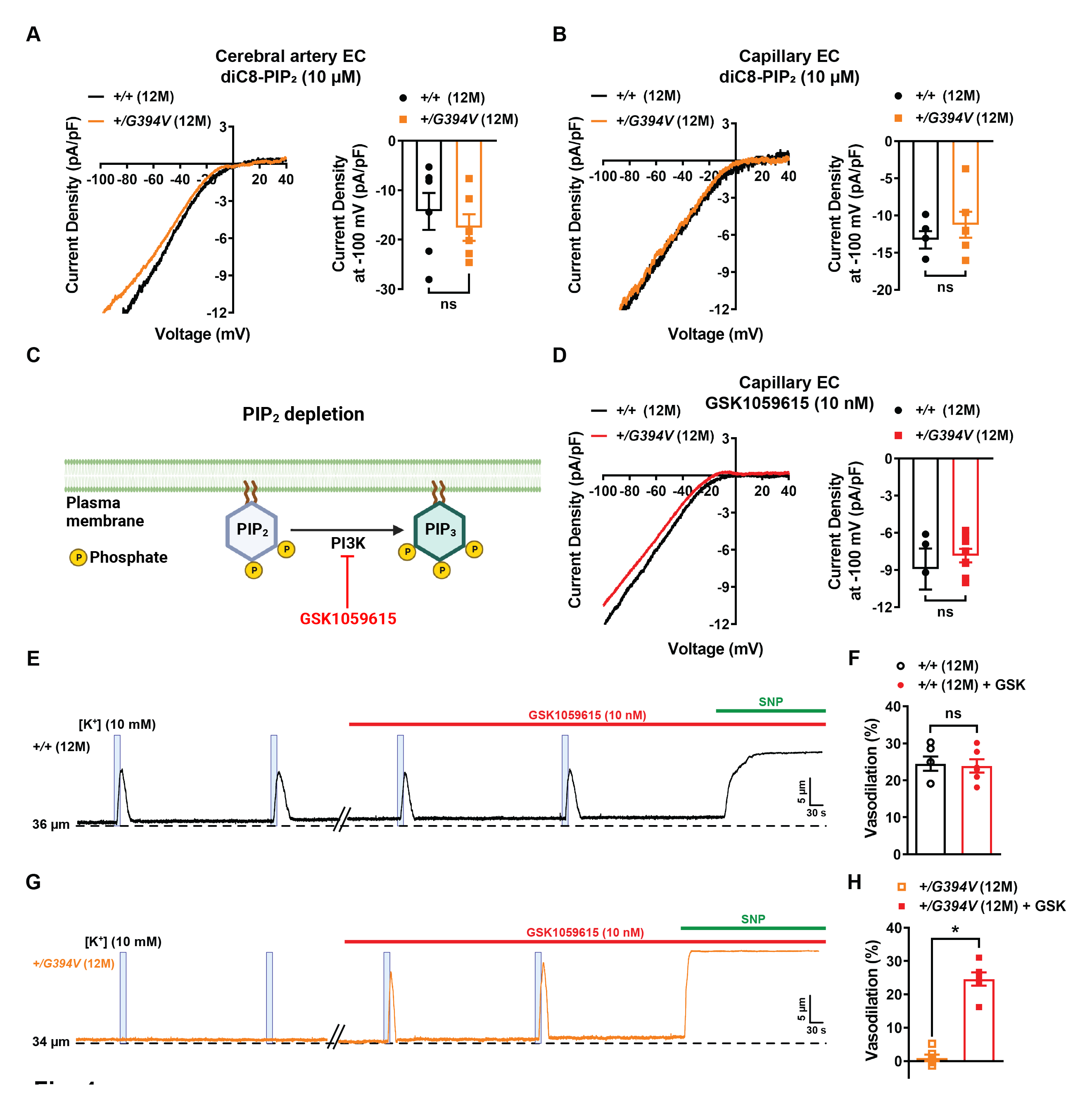
PIP_2_ depletion reduces Kir2.1 currents in 12 M-old Col4a1^+/G394V^ mice. (A) Representative I-V traces and summary data showing Kir2.1 current densities in freshly isolated cerebral artery ECs from 12 M-old *Col4a1^+/+^* and *Col4a1^+/G394V^* mice with internal solution supplemented with diC8-PIP2 (10 µM) (n = 6 cells from 4 animals per group, ns = not significant, unpaired t-test). (B) Representative I-V traces and summary data showing Kir2.1 current densities in freshly isolated brain capillary ECs from 12 M-old *Col4a1^+/+^* and *Col4a1^+/G394V^* mice with internal solution supplemented with diC8-PIP2 (10 µM) (n = 5–6 cells from 4 to 5 animals per group, ns = not significant, unpaired t-test). (C) PIP_2_ depletion pathway. (D) Representative I-V traces and summary data showing Kir2.1 current densities in freshly isolated brain capillary ECs from 12 M-old *Col4a1^+/+^* and *Col4a1^+/G394V^* mice treated with the PI3K blocker GSK1059615 (10 nM) (n = 4–8 cells from 3 to 6 animals per group, ns = not significant, unpaired t-test). (E and F) Representative trace (E) and summary data (F) showing K^+^ (10 mM, blue box)-induced dilation of upstream arterioles in preparations from 12 M-old *Col4a1^+/+^* mice before and after superfusing the PI3K blocker GSK1059615 (10 nM, 30 min) (n = 6 preparations from 5 animals per group, ns = not significant, paired t-test). (G and H) Representative trace (G) and summary data (H) showing K^+^ (10 mM, blue box)-induced dilation of upstream arterioles in preparations from 12 M-old *Col4a1^+/G394V^* mice before and after superfusing the PI3K blocker GSK1059615 (10 nM, 30 min) (n = 6 preparations from 5 animals per group, *p<0.05, paired t-test).

To determine if elevated PI3K activity is also responsible for cerebral microvascular dysfunction in mutant mice, capillary-to-arteriole dilation was assessed before and after treatment with GSK1059615 (10 nM, 30 min). This treatment had no effect on preparations from 12 M-old control animals (Fig. 4E and F). In contrast, PI3K block fully restored K^+^-induced dilation of upstream arterioles from 12 M-old *Col4a1^+/G394V^* mice (Fig. 4G and H). In control studies, we found that the vehicle for GSK1059615 (0.01% v/v DMSO) did not affect capillary-to-arteriole dilation (Fig. 4 – supplement 2A to D) or maximal vasodilation in response to SNP (Fig. 4 – supplement 2E and F). These data suggest that elevated PI3K activity is responsible for the loss of capillary-mediated dilation in *Col4a1* mutant mice.

### Chronic PI3K inhibition restores Kir2.1 currents, K^+^-induced dilation, functional hyperemia, and memory deficits in 12 M-old Col4a1^+/G394V^ mice

To determine if chronic inhibition of PI3K could resolve deficits in functional hyperemia and memory in mutant mice, 12 M-old *Col4a1^+/G394V^* and control mice were injected subcutaneously with GSK1059615 (10 mg/kg) or vehicle each day for 28 days (Fig. 5A). At the end of the treatment, we found that Kir2.1 currents in brain capillary ECs from *Col4a1^+/G394V^* mice injected with GSK1059615 were fully restored to control levels, whereas vehicle treatment had no effect (Fig. 5B). GSK1059615 treatment fully restored capillary-to-arteriole dilation in *ex vivo* microvascular preparations from *Col4a1^+/G394V^* mice (Fig. 5C and D). These data demonstrate that defects in brain capillary EC Kir2.1 channel function and associated microvascular vasomotor dysfunction can be restored by chronic PI3K blockade.

**Fig. 5.**
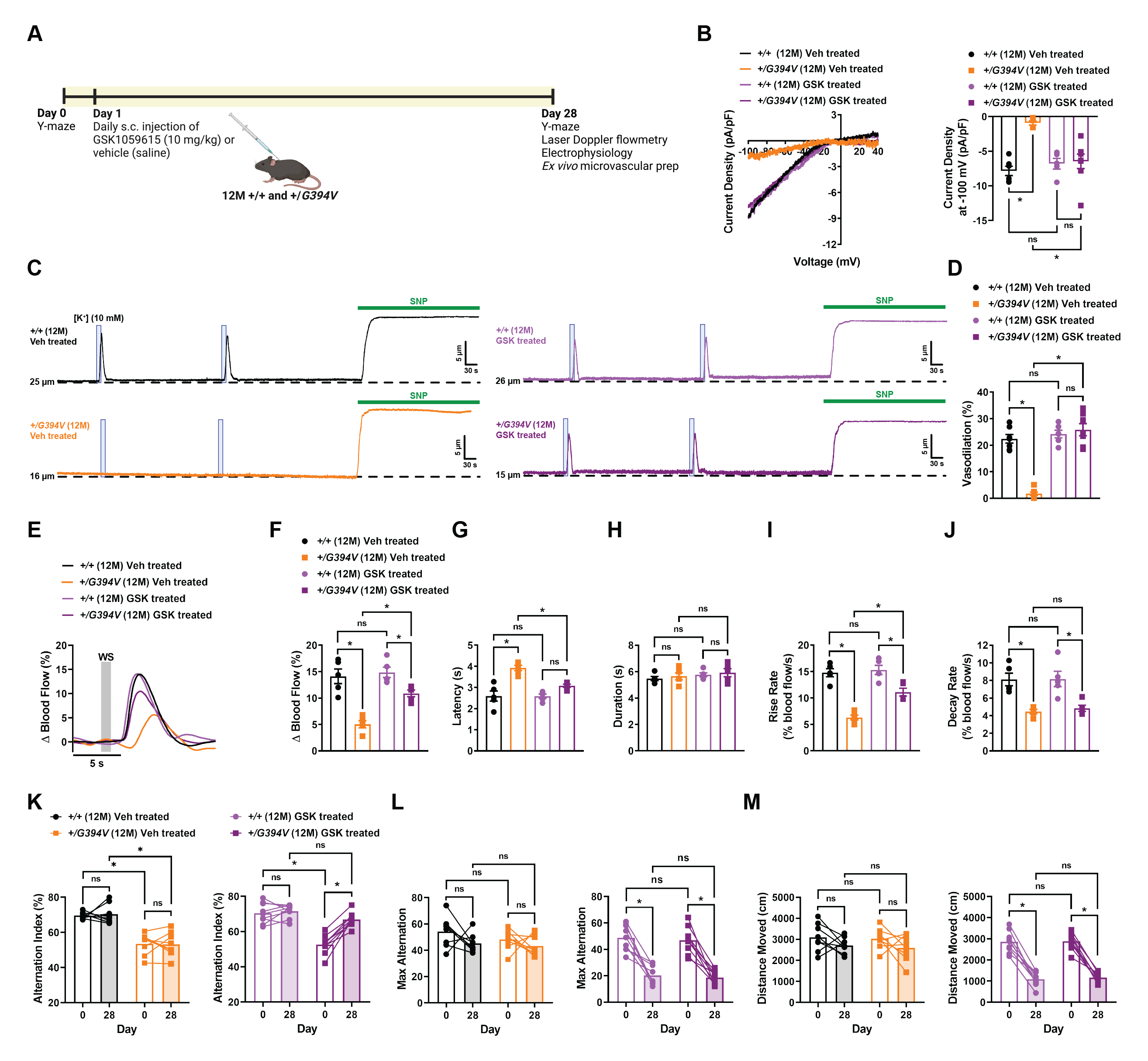
Chronic PI3K inhibition restores Kir2.1 currents, K^+^-induced dilation, functional hyperemia, and memory deficits in 12 M-old Col4a1^+/G394V^ mice. (A) Illustration showing GSK1059615 treatment plan. (B) Representative I-V traces and summary data showing Kir2.1 current densities in freshly isolated capillary ECs from 12 M-old Col4a1^+/+^ and Col4a1^+/G394V^ mice treated with vehicle (saline) or GSK1059615 (10 mg/kg) for 28 days, s.c. (n = 5–8 cells from 3 to 4 animals per group, *p<0.05, ns = not significant, non-repeated measures two-way ANOVA). (C and D) Representative traces (C) and summary data (D) showing K^+^ (10 mM, blue box)-induced dilation of upstream arterioles in preparations from 12 M-old Col4a1^+/+^ and Col4a1^+/G394V^ mice treated with vehicle (saline) or GSK1059615 (10 mg/kg) for 28 days, s.c. (n = 6–7 preparations from 3 animals per group, *p<0.05, ns = not significant, non-repeated measures two-way ANOVA). (E and F) Representative traces (E) and summary data (F) showing the increase in blood flow following 1 s contralateral whisker stimulation (WS) in 12 M-old Col4a1^+/+^ and Col4a1^+/G394V^ mice treated with vehicle (saline) or GSK1059615 (10 mg/kg) for 28 days, s.c. (n = 5 animals per group, *p<0.05, ns = not significant, non-repeated measures two-way ANOVA). (G to J) Latency (G), duration (H), rise rate (I), and decay rate (J) were also analyzed (n= 5 animals per group, *p<0.05, ns = not significant, non-repeated measures two-way ANOVA). (K) Summary data showing alternation index, an indicator spatial working memory, of 12 M-old Col4a1^+/+^ and Col4a1^+/G394V^ mice before and after treatment with vehicle (saline) or GSK1059615 (10 mg/kg) for 28 days, s.c. (n = 8–9 animals per group, *p<0.05, ns = not significant, repeated measures two-way ANOVA). (L and M) Summary data showing max alternation (L) and distance moved (M), indicative of exploratory activity, in 12 M-old Col4a1^+/+^ and Col4a1^+/G394V^ mice before and after treatment with vehicle (saline) or GSK1059615 (10 mg/kg) for 28 days, s.c. (n = 8–9 animals per group, *p<0.05, ns = not significant, repeated measures two-way ANOVA).

GSK1059615 treatment also increased the magnitude of blood flow increases in the somatosensory cortex induced whisker stimulation (1 s) in *Col4a1^+/G394V^* mice compared to vehicle-treated animals (Fig. 5E and F). The latency and rise rates of the blood flow response were also improved (Fig. 5G to J). Similar outcomes were observed for 2 s and 5 s stimulation protocols (Fig. 5 – supplement 1). In control studies, stimulation of ipsilateral whiskers failed to produce any change in the blood flow (Fig. 5 – supplement 2). These data demonstrate that chronic PI3K inhibition can restore functional hyperemia in *Col4a1^+/G394V^* mice.

In a longitudinal study, we found that GSK1059615 treatment significantly improved the Y-maze spontaneous alternation behavior of 12 M-old *Col4a1^+/G394V,^* whereases vehicle-treated mutant mice did not improve. The performance of GSK1059615-treated *Col4a1^+/G394V^* mice did not differ from that of age-matched control animals (Fig. 5K). Interestingly, GSK1059615 treatment reduced the maximum number of alternations (Fig. 5L) and the total distance moved (Fig. 5M) for control and mutant mice. This unexpected side effect of prolonged GSK1059615 administration may be related to fatigue symptoms reported by cancer patients treated with PI3K inhibitors ^46^. Despite this potential limitation, our data identify PI3K inhibition as a novel therapeutic strategy for treating cognitive impairment associated with some forms of cSVD.

## Discussion

Brain injuries and loss of fundamental blood flow control mechanisms due to cSVDs are significant causes of adult dementia. The brain pathology of *Col4a1^+/G394V^* cSVD mice used in this investigation was mild compared to other models, but NVC and functional hyperemia were significantly impaired. Thus, these animals allow the effects of impaired vascular control to be investigated independently of severe brain damage. Aged mutants performed worse than control mice in behavioral tests of working and recognition memory, suggesting that defects in cerebral blood flow control mechanisms are a primary cause of cognitive impairment in these animals. The fundamental defect leading to impaired NVC was the loss of Kir2.1 channel activity in brain capillary and arterial ECs due to PIP_2_ depletion. Accordingly, chronic PI3K blockade rescued Kir2.1 currents, NVC, functional hyperemia, and improved memory function in mutant mice, providing evidence that cognitive impairment associated with cSVD can be resolved by improving blood flow regulation in the brain.

Missense mutations in *COL4A1* and *COL4A2* that alter G residues in collagen triple helices are the most common cause of Gould syndrome ^47-49^. Such mutations prevent the proper assembly of collagen α1α1α2(IV) heterotrimers, potentially leading to intracellular retention, ER stress, and/or disrupting basement membranes to cause cSVD and other pathologies ^50^. The impact of specific disease-causing point mutations is highly variable and position dependent. Notably, spontaneous ICH is less severe for mutations nearer to the amino terminus, such as the *Col4a1^G394V^* mutation, and more severe for mutations closer to the carboxyl terminus ^31,51^. Consistent with this finding, *Col4a1^+/G394V^* mice did not show overt cerebral pathology in MRI scans. However, capillary-mediated NVC and functional hyperemia were impaired in an age-dependent manner. The age dependence of this pathology may be related to the unique properties of collagen. Collagens are extraordinarily durable - the *in vivo* half-life of collagen I, present in ligaments and tendons, is more than 100 years ^52^. The turnover rate of collagen α1α1α2(IV) is not precisely known, but an early study reported that the half-life of “vascular collagen” in rat aorta and mesenteric arteries was ∼70 days ^53^. Further, collagen synthesis peaks during late embryonic development and growth as basement membranes form and then markedly decreases with age ^54,55^. Our data suggest that *Col4a1^+/G394V^* mice successfully develop and maintain functional basement membranes early in life when collagen production is maximal. We propose that diminishing collagen production and increasing degradation during aging result in a slow diminution of basement membranes over time, eventually leading to defects by middle age. It is conceivable that a similar process could contribute to idiopathic forms of age-related cSVD in humans.

Elevated TGF-β signaling has been recently described in *Col4a1* mutant mice. Genetic knockdown of TGF-β ligand and blocking TGF-β signaling using neutralizing antibodies or pharmacological inhibitors improved ocular and cerebrovascular pathogenesis associated with *Col4a1* mutations^32,56,57^. Activation of TGF-β receptors can stimulate PI3K activity through a non-canonical signaling pathway involving the TRAF6 ubiquitin ligase ^58,59^, suggesting that enhanced TGF-β signaling contributes to PIP_2_ insufficiency in *Col4a1^+/G394V^* mice. Further, disinhibition of TGF-β signaling causes CARASIL (cerebral autosomal recessive arteriopathy with subcortical infarcts and leukoencephalopathy), a rare genetic form of cSVD ^60^, suggesting that disturbances in the TGF-β pathway may be a generalized mechanism of cerebral vascular dysfunction.

Altered PIP_2_ metabolism has emerged as a central pathogenic mechanism in multiple forms of cerebral microvascular disease. Prior reports show that PIP_2_ insufficiency diminishes Kir2.1 channel activity in brain capillary ECs in mouse models of CADASIL cSVD ^22^ and familial Alzheimer’s disease ^41^. The current data show that the same defect underlies impaired functional hyperemia and memory deficits in *Col4a1^+/G394V^* mutant mice. Prior studies provide evidence that reduced levels of ATP in brain capillary ECs diminished the production of PIP_2_ ^22^, whereases our findings indicate that increased activity of PI3K decreased PIP_2_ bioavailability in *Col4a1^+/G394V^* mice. We also show that PIP_2_ levels and Kir2.1 channel activity are diminished in cerebral arteriolar ECs from 12 M-old *Col4a1* mutant mice. In contrast, these cells are unaffected in CADASIL and 5xFAD animals, demonstrating cellular heterogeneity of PIP_2_ depletion mechanisms among the different disease models. Despite these differences, we propose that repairing defective NVC by increasing PIP_2_ levels in brain capillary ECs is a viable therapeutic strategy for multiple forms of cerebrovascular disease. We provide proof-of-concept by targeting the PI3K pathway to preserve PIP_2_, thereby restoring functional hyperemia and improving memory function in *Col4a1* mutant mice. Several types of PI3K inhibitors are approved by the FDA for use against advanced cancers ^46^ and could rapidly advance to clinical trials for the treatment of cSVDs.

## Materials and methods

### Chemical and reagents

Chemicals and other reagents were obtained from Sigma-Aldrich, Inc. (St. Louis, MO, USA) unless otherwise specified.

### Animals

Young adult (3 M-old) and middle-aged (12 M-old) male and female littermate *Col4a1^+/+^* and *Col4a1^+/G394V^* mice were used in this study ^25,61^. Animals were maintained in individually ventilated cages (<5 mice/cage) with *ad libitum* access to food and water in a room with controlled 12-hour light and dark cycles. All animal care procedures and experimental protocols involving animals complied with the NIH *Guide for the Care and Use of Laboratory Animals* and were approved by the Institutional Animal Care and Use Committees at the University of Nevada, Reno, and the University of California, San Francisco. Arteries for *ex vivo* experimentation were harvested from mice anesthetized with isoflurane (Baxter Healthcare, Deerfield, IL, USA) and euthanized by decapitation and exsanguination. Brains were isolated and placed in ice-cold Ca^2+^-free physiological saline solution (Mg-PSS; 134 mM NaCl, 5 mM KCl, 2 mM MgCl_2_, 10 mM HEPES, 10 mM glucose, 0.5% bovine serum albumin, pH 7.4 with NaOH).

### In vivo magnetic resonance imaging

All *in vivo* magnetic resonance (MR) experiments were conducted on a 14.1 Tesla vertical MR system (Agilent Technologies, Palo-Alto, CA, USA) equipped with 100G/cm gradients and a single tuned millipede ^1^H proton coil (ØI = 40mm). For each imaging session, mice were anesthetized using isoflurane (1-1.5% in O2) and positioned in a dedicated cradle maintaining constant anesthesia and placed in the MR bore; respiration and temperature were continuously monitored during all acquisitions to ensure animal well-being and data reproducibility. Susceptibility Weighted Imaging (SWI) was performed to detect the potential presence of hemorrhagic lesions, using the following parameters: gradient-echo scheme, field of view (FOV) = 20 x 20 mm^2^, matrix = 256 x 256, 16 slices, 0.4 mm slice thickness, 0.1 mm interslice gap, number of averages = 16, echo time (TE)/ repetition time (TR) = 4.60 / 140 ms, flip angle = 10 degrees. T2-weighted (T2W) images were also acquired in a fast-spin-echo scheme to calculate brain and ventricle volumes, using the same FOV geometry as SWI and the following parameters: number of averages = 8, TE/TR = 21.38 / 2500 ms, flip angle = 90 degrees. For each animal, total brain and ventricles were manually delineated on the T2W images, their volumes calculated, and the ventricle/brain ratio computed.

### Isolation of native brain capillary ECs

Individual brain capillary ECs were obtained as previously described ^18,19^. Brains were denuded of surface vessels, and two 1 mm-thick were excised and homogenized in artificial cerebrospinal fluid (aCSF; 124 mM NaCl, 3 mM KCl, 2 mM MgCl_2_, 2 mM CaCl_2_, 1.25 mM NaH_2_PO_4_, 26 mM NaHCO_3_, and 4 mM glucose) using a Dounce homogenizer. The homogenate was filtered through a 70 µM filter, and capillary networks captured on the filter were transferred to a new tube. Individual cells were isolated by enzymatic digestion with 0.5 mg/ml neutral protease and 0.5 mg/ml elastase (Worthington Biochemical Corporation) in endothelial cell (EC) isolation solution (55 mM NaCl, 80 mM Na-glutamate, 6 mM KCl, 2 mM MgCl_2_, 10 mM glucose, 0.1 mM CaCl_2_,10 mM HEPES, pH 7.3) for 45 min at 37°C. Following this, 0.5 mg/ml collagenase type I (Worthington Biochemical Corporation) was added, and a second 2-min incubation at 37°C was performed. Digested networks were washed in ice-cold EC isolation solution, then triturated with a fire-polished glass Pasteur pipette to produce individual ECs. All cells used for this study were freshly dissociated on the day of experimentation.

### Isolation of cerebral arteriolar ECs

Single arterial ECs were isolated as previously described ^18^. Cerebral arteries were dissected from mouse brains and washed in Mg-PSS. Arteries were transferred to EC isolation solution supplemented with 0.5 mg/mL neutral protease and 0.5 mg/mL elastase (Worthington Biochemical Corporation, Lakewood, NJ, USA), and incubated for 40 min at 37°C. Following this, 0.5 mg/mL collagenase type I (Worthington Biochemical Corporation) was added for an additional 2 min incubation at 37°C. A single-cell suspension was prepared by washing digested arteries three times with EC isolation solution to remove enzymes and triturating with a fire-polished glass pipette to dissociate cells. All cells used for this study were freshly dissociated on the day of experimentation.

### Whole-cell patch-clamp electrophysiology

Enzymatically isolated native ECs were transferred to a recording chamber (Warner Instruments, Hamden, CT, USA) and allowed to adhere to glass coverslips for 15 minutes at room temperature. Pipettes were fabricated from borosilicate glass (1.5 mm outer diameter, 1.17 mm inner diameter; Sutter Instruments, Novato, CA, USA), fire-polished to yield a tip resistance of 3–6 MΩ. Currents were recorded at room temperature using an Axopatch 200B amplifier (Molecular Devices, San Jose, CA, USA) equipped with an Axon CV 203BU headstage and Digidata 1440A digitizer (Molecular Devices) for all patch-clamp electrophysiology experiments. Currents were filtered at 1 kHz and digitized at 10 kHz. Kir2.1 currents were recorded using the conventional whole-cell configuration at a holding potential of -50 mV, with 400 ms ramps from -100 to +40 mV. The external bathing solution was composed of 134 mM NaCl, 6 mM KCl, 1 mM MgCl_2_, 2 mM CaCl_2_, 10 mM glucose, and 10 mM HEPES. The composition of the pipette solution was 10 mM NaCl, 30 mM KCl, 10 mM HEPES, 110 mM K^+^ aspartate, and 1 mM MgCl_2_ (pH 7.2). Kir2.1 currents were activated by increasing extracellular [K^+^] concentration from 6 to 60 mM ([NaCl] adjusted to maintain isotonic solution) and blocked using BaCl_2_ (10 µM). All recordings were performed at room temperature. Clampex and Clampfit software (pClamp version 10.2; Molecular Devices) were used for data acquisition and analysis, respectively.

### Pressure myography

The current best practices guidelines for pressure myography experiments were followed ^30^. Pressure myograph experiments were performed on cerebral arteries and parenchymal arteriole-capillary preparations ^19^.

Surface cerebral arteries were carefully isolated and mounted between two glass cannulas (approximate outer diameter 40–50 μm) in a pressure myograph chamber (Living Systems Instrumentation, St Albans City, VT, USA) and secured by a nylon thread. Intraluminal pressure was controlled using a servo-controlled peristaltic pump (Living Systems Instrumentation), and preparations were visualized with an inverted microscope (Accu-Scope Inc., Commack, NY, USA) coupled to a USB camera (The Imaging Source LLC, Charlotte, NC, USA). Changes in luminal diameter were assessed using IonWizard software (version 7.2.7.138; IonOptix LLC, Westwood, MA, USA). Arteries were bathed in warmed (37°C), oxygenated (21% O_2_, 6% CO_2_, 73% N_2_) PSS (119 mM NaCl, 4.7 mM KCl, 21 mM NaHCO_3_, 1.17 mM MgSO_4_, 1.8 mM CaCl_2_, 1.18 mM KH_2_PO_4_, 5 mM glucose, 0.03 mM EDTA) at an intraluminal pressure of 5 mmHg. Following equilibration for 15 min, intraluminal pressure was increased to 110 mmHg, and vessels were stretched to their approximate *in vivo* length, after which pressure was reduced back to 5 mmHg for an additional 15 min. Vessel viability was assessed for each preparation by evaluating vasoconstrictor responses to high extracellular [K^+^] PSS, made isotonic by adjusting the [NaCl] (60 mM KCl, 63.7 mM NaCl). Arteries that showed less than 10% constriction in response to elevated extracellular [K^+^] were excluded from further investigation. Changes in lumen diameter were recorded at different concentrations of extracellular [K^+^] (8 to 60 mM, [NaCl] adjusted to maintain isotonic solution). Arteries were pressurized to 20 mmHg and superfused with 10 nM endothelin-1 to induce vasoconstriction. Passive lumen diameter was determined by superfusing vessels Ca^2+^-free PSS supplemented with EGTA (2 mM) and the voltage-dependent Ca^2+^ channel blocker diltiazem (10 μM) to inhibit SMC contraction. Change in diameter was calculated at each [K^+^] concentration as the change in diameter (%) = (change in lumen diameter / passive diameter) × 100.

Parenchymal arterioles with intact capillary segments deriving from the middle cerebral artery were carefully dissected, cannulated, and secured onto a pressure myograph chamber using the same method described above ^19^. Preparations were bathed in a warmed, oxygenated aCSF solution at an intraluminal pressure of 5 mmHg. Following equilibration for 15 min, intraluminal pressure was increased to 20 mmHg and superfused with 10 nM endothelin-1 to induce vasoconstriction. Localized application of drugs onto the capillary extremities was achieved by placing a micropipette attached to a Picospritzer III (Parker Hannifin, Cleveland, OH, USA) adjacent to capillary segments. Capillaries were stimulated by locally applying a 7 s pulse of aCSF containing elevated [K^+^] (10 mM, [NaCl] adjusted to maintain isotonic solution) onto capillary extremities. To determine preparation viability, SNP (10 µM) was superfused into the circulating aCSF. Changes in lumen diameter were calculated as vasodilation (%) = (change in lumen diameter / baseline diameter) × 100.

### Assessment of functional hyperemia in the brain using laser Doppler flowmetry

Functional hyperemia in the brain was assessed as previously described ^19^. Mice were anesthetized with isoflurane (5% induction, 2% maintenance), the head was immobilized in a stereotaxic frame, and the skull was exposed. The skull of the right hemisphere was carefully thinned using a drill to visualize the surface vasculature of the somatosensory cortex. Isoflurane anesthesia was replaced with combined α-chloralose (50 mg/kg, i.p.) and urethane (750 mg/kg, i.p.) to eliminate confounding vasodilatory effects of isoflurane. Changes in perfusion were assessed via a laser-Doppler flowmetry probe (PeriFlux System PF5000, Perimed AB, Jakobsberg, Sweden) positioned directly above the somatosensory cortex. The contralateral whiskers were stimulated for either 1, 2, or 5 s, and changes in perfusion were recorded. Contralateral whiskers were stimulated three times at 2 min intervals. Ipsilateral whiskers were also stimulated as a control for potential vibration artifacts. Changes in perfusion were calculated as %Δ Blood flow = (perfusion during stimulus / baseline perfusion) × 100. Kinetics of the response, including latency, duration, rise rate, and decay rate, were also obtained and analyzed.

### Y-Maze behavioral assay

Memory function was assessed using the Y-maze (Maze Engineers, Skokie, IL, USA) using two different configurations.

The spontaneous alternation behavioral assay was used to assess short-term spatial working memory ^36^. Mice were placed into one of the three arms of the maze (start arm) and were allowed to explore all three arms for 10 minutes. Session videos were recorded and analyzed using Ethovision XT software (version 16.0.1536, Noldus Information Technology, Leesburg, VA, USA). Spontaneous alternation was evaluated by scoring the order of entries into each arm during the 10-minute recording period. Spontaneous alternation was calculated as alternation index (%) = (number of spontaneous alternations/max alternation) × 100, where the spontaneous alternation is defined as the number of consecutive entries into each arm of the maze in any order without any repeats, and the max alternation is the total number of alternations possible (max alternation = total number of arm entries – 2).

The novel arm configuration was used to evaluate longer-term spatial reference memory ^36^. This test was performed across 2 days. On day one, mice were placed into one of the three arms of the maze (start arm) and allowed to explore only two arms for 10 minutes (training trial). On day two, the test trial was conducted with the closed arm opened, which served as the novel arm. Mice were returned to the maze via the same start arm and were allowed to explore all three arms for 10 minutes. Session videos were recorded and analyzed using Ethovision XT software, and the time spent in the novel arm and entries into the novel arm were measured and analyzed.

### Measurement of [ATP] in brain capillary ECs

The intracellular [ATP] was determined by lysing brain capillary ECs and quantifying luminescence produced by luciferase-induced conversion of ATP to light using the CellTiter-Glo Assay 3D (Promega, Madison, WI, USA). Approximately ∼500 ECs were added to each reaction. To further ensure equal input of ECs, the CellTiter-Glo 3D Assay was multiplexed with CellTox Green Assay (Promega), a fluorescent dye that selectively and quantitatively binds double-stranded DNA. Dye fluorescence is directly proportional to DNA concentration and the number of cells in each assay. ATP concentration (luminescence) was normalized to the number of cells (fluorescence) to determine intracellular ATP per cell. Luminescence and fluorescence (485_Ex_/538_Em_) were measured using a FlexStation 3 (Molecular Devices). A serial tenfold dilution of ATP (1 nM to 1 uM; 80 µl contains 8^-14^ to 8^-11^ moles of ATP, respectively) was measured to calibrate the linear working range of ATP detection. Data were expressed as the ratio of ATP luminescence/DNA fluorescence.

### Statistical analysis

All summary data are presented as means ± SEM. Statistical analyses and graphical presentations were performed using GraphPad Prism software (version 9.4.1, GraphPad Software, Inc, USA). The value of n refers to the number of cells for patch-clamp electrophysiology experiments and measurement of [ATP], vessel preparations for myography experiments, and animals used for MRI acquisition, the Y-maze behavioral assays and functional hyperemia assessment. Statistical analyses were performed using Student’s paired or unpaired two-tailed t-test, or repeated measures or non-repeated measures two-way analysis of variance (ANOVA) with a Šidák correction for multiple comparisons. A value of p<0.05 was considered statistically significant.

## Author contributions

S.E. and D.B.G. initiated and supervised the project. S.E. designed the experiments. X.G. and M.M.C. performed *in vivo* MRI experiments. S.A., E.Y., and A.S.S. performed patch-clamp electrophysiology experiments. E.Y. and P.T. performed myography experiments. P.T. conducted *in vivo* functional hyperemia experiments and performed Y-maze behavioral tests. E.Y. conducted luciferase assay for the measurement of the cellular level of ATP. P.T., E.Y., S.A., and S.E. analyzed the data. P.T. and S.E. wrote the manuscript and prepared the figures. P.T., E.Y., C.L.D., D.B.G., and S.E. revised the manuscript.

## Data and materials availability

All data needed to evaluate the conclusions are present in the paper or the Supplementary Materials.

## Supplemental figure legends

***Fig. 1 – supplement 1. Kir2.1 currents in cerebral artery ECs.*** (A) Illustration of the cerebral artery EC isolation procedure. Scale bar = 10 µm. (B) Representative I-V traces and summary data showing Kir2.1 current densities in freshly isolated cerebral artery ECs from 3 M-old *Col4a1^+/+^* and *Col4a1^+/G394V^* mice (n = 5–6 cells from 5 animals per group, ns = not significant, unpaired t-test). (C) Representative I-V traces and summary data showing Kir2.1 current densities in freshly isolated cerebral artery ECs from 12 M-old *Col4a1^+/+^* and *Col4a1^+/G394V^* mice (n = 6–10 cells from 3 to 4 animals per group; *p<0.05, unpaired t-test). (D and E) Representative traces (D) and summary data (E) demonstrating the vasodilator response to raising extracellular [K^+^] in cerebral arteries from 12 M-old *Col4a1^+/+^* and *Col4a1^+/G394V^* mice (n = 9–13 preparations from 6 to 10 animals per group, *p<0.05, non-repeated measures two-way ANOVA). (F) Summary data showing the contractile response to 30 and 60 mM [K^+^] in cerebral arteries from 12 M-old *Col4a1^+/+^* and *Col4a1^+/G394V^* mice (n = 7–12 preparations from 5 to 10 animals per group, ns = not significant, unpaired t-test).

***Fig. 1 – supplement 2. High field magnetic resonance imaging.*** (A) Representative susceptibility weighted (SWI) and T2 weighted (T2W) MRI images from 12 M-old *Col4a1^+/+^* and *Col4a1^+/G394V^* mice. (B) Quantification of ventricle/brain ratio for 12 M-old *Col4a1^+/+^* and *Col4a1^+/G394V^* mice (n = 7 animals per group, ns = not significant, unpaired t-test).

***Fig. 2 – supplement 1. Functional hyperemic response following 2 s whisker stimulation.*** (A and B) Representative traces (A) and summary data (B) showing the increase in blood flow following 2 s contralateral whisker stimulation (WS) in 3 M-old *Col4a1^+/+^* and *Col4a1^+/G394V^* mice (n = 6 animals per group, ns = not significant, unpaired t-test). (C to F) Latency (C), duration (D), rise rate (E) and decay rate (F) were also analyzed (n= 6 animals per group, ns = not significant, unpaired t-test). (G and H) Representative traces (G) and summary data (H) showing the increase in blood flow following 2 s contralateral WS in 12 M-old *Col4a1^+/+^* and *Col4a1^+/G394V^* mice (n = 6 animals per group, *p<0.05, unpaired t-test). (I to L) Latency (I), duration (J), rise rate (K) and decay rate (L) were also analyzed (n = 6 animals per group, *p<0.05, ns = not significant, unpaired t-test).

***Fig. 2 – supplement 2. Ipsilateral whisker stimulation.*** (A to C) Summary data showing no change in blood flow following ipsilateral whisker stimulation for 1 s (A), 2 s (B), and 5 s (C) in 3 M- and 12 M-old *Col4a1^+/+^* and *Col4a1^+/G394V^* mice (n = 6 animals per group, ns = not significant, unpaired t-test).

***Fig. 4 – supplement 1. ATP levels in brain capillary ECs.*** (A) PIP_2_ synthesis pathway. (B) Summary data showing normalized [ATP] in brain capillary ECs from 12 M-old *Col4a1^+/+^* and *Col4a1^+/G394V^* mice. (n = 9 sets of ∼500 cells from three mice per group, ns = not significant, unpaired t-test). (C) PIP_2_ depletion pathway. (D) Representative I-V traces and summary data showing Kir2.1 current densities in freshly isolated brain capillary ECs treated with the PLC inhibitor U73122 (10 µM) from 12 M-old *Col4a1^+/+^* and *Col4a1^+/G394V^* mice (n = 4 cells from 3 animals per group, *p<0.05, unpaired t-test).

***Fig. 4 – supplement 2. GSK1059615 vehicle and SNP response.*** (A and B) Representative trace (A) and summary data (B) showing K^+^ (10 mM, blue box)-induced dilation of upstream arterioles in preparations before and after superfusing the vehicle (0.01% v/v DMSO, 30 min) from 12 M-old *Col4a1^+/+^* mice (n = 6 preparations from 4 animals per group, ns = not significant, paired t-test). (C and D) Representative trace (C) and summary data (D) showing K^+^ (10 mM, blue box)-induced dilation of upstream arterioles in preparations before and after superfusing the vehicle (0.01% v/v DMSO, 30 min) from 12 M-old *Col4a1^+/G394V^* mice (n = 6 preparations from 4 animals per group, ns = not significant, paired t-test). (E and F) Summary data showing that the dilation produced by superfusing SNP (10 µM) in the presence of GSK1059615 (10 nM, 30 min) (E), and vehicle (0.01% v/v DMSO, 30 min) (F) in preparations from 12 M-old *Col4a1^+/+^* and *Col4a1^+/G394V^* mice (n = 6 preparations from 3 to 5 animals per group, ns = not significant, unpaired t-test).

***Fig. 5 – supplement 1. Functional hyperemic response following 2 s and 5 s whisker stimulation in GSK1059615-treated animals.*** (A and B) Representative traces (A) and summary data (B) showing the increase in blood flow following 2 s contralateral whisker stimulation (WS) in 12 M-old *Col4a1^+/+^* and *Col4a1^+/G394V^* mice treated with vehicle (saline) or GSK1059615 (10 mg/kg) for 28 days, s.c. (n = 5 animals per group, *p<0.05, ns = not significant, non-repeated measures two-way ANOVA). (C to F) Latency (C), duration (D), rise rate (E) and decay rate (F) were also analyzed (n= 5 animals per group, *p<0.05, ns = not significant, non-repeated measures two-way ANOVA). (G and H) Representative traces (G) and summary data (H) showing the increase in blood flow following 5 s contralateral WS in 12 M-old *Col4a1^+/+^* and *Col4a1^+/G394V^* mice treated with vehicle (saline) or GSK1059615 (10 mg/kg) for 28 days, s.c. (n = 5 animals per group, *p<0.05, ns = not significant, non-repeated measures two-way ANOVA). (I to L) Latency (I), duration (J), rise rate (K) and decay rate (L) were also analyzed (n= 5 animals per group, *p<0.05, ns = not significant, non-repeated measures two-way ANOVA).

***Fig. 5 – supplement 2. Ipsilateral whisker stimulation in GSK1059615 treated animals.*** (A to C) Summary data showing no change in blood flow following ipsilateral whisker stimulation for 1 s (A), 2 s (B), and 5 s (C) in 12 M-old *Col4a1^+/+^* and *Col4a1^+/G394V^* mice treated with vehicle (saline) or GSK1059615 (10 mg/kg) for 28 days, (n = 5 animals per group, ns = not significant, non-repeated measures two-way ANOVA).

## References

1 Corriveau, R. A. et al. The Science of Vascular Contributions to Cognitive Impairment and Dementia (VCID): A Framework for Advancing Research Priorities in the Cerebrovascular Biology of Cognitive Decline. Cell Mol Neurobiol 36, 281–288 (2016). https://doi.org:10.1007/s10571-016-0334-7

2 Wardlaw, J. M., Smith, C. & Dichgans, M. Mechanisms of sporadic cerebral small vessel disease: insights from neuroimaging. Lancet Neurol 12, 483–497 (2013). https://doi.org:Doi10.1016/S1474-4422(13)70060-7

3 Debette, S., Schilling, S., Duperron, M. G., Larsson, S. C. & Markus, H. S. Clinical Significance of Magnetic Resonance Imaging Markers of Vascular Brain Injury A Systematic Review and Meta-analysis. Jama Neurol 76, 81–94 (2019). https://doi.org:10.1001/jamaneurol.2018.3122

4 Dichgans, M. & Leys, D. Vascular Cognitive Impairment. Circ Res 120, 573–591 (2017). https://doi.org:10.1161/CIRCRESAHA.116.308426

5 Shi, Y. & Wardlaw, J. M. Update on cerebral small vessel disease: a dynamic whole-brain disease. Stroke Vasc Neurol 1, 83–92 (2016). https://doi.org:10.1136/svn-2016-000035

6 Evans, L. E. et al. Cardiovascular comorbidities, inflammation, and cerebral small vessel disease. Cardiovasc Res 117, 2575–2588 (2021). https://doi.org:10.1093/cvr/cvab284

7 van Dinther, M. et al. Extracerebral microvascular dysfunction is related to brain MRI markers of cerebral small vessel disease: The Maastricht Study. Geroscience 44, 147–157 (2022). https://doi.org:10.1007/s11357-021-00493-0

8 Jokumsen-Cabral, A., Aires, A., Ferreira, S., Azevedo, E. & Castro, P. Primary involvement of neurovascular coupling in cerebral autosomal-dominant arteriopathy with subcortical infarcts and leukoencephalopathy. J Neurol 266, 1782–1788 (2019). https://doi.org:10.1007/s00415-019-09331-y

9 Roy, C. S. & Sherrington, C. S. On the Regulation of the Blood-supply of the Brain. J Physiol 11, 85–158 117 (1890).

10 Jeanne, M. & Gould, D. B. Genotype-phenotype correlations in pathology caused by collagen type IV alpha 1 and 2 mutations. Matrix Biol 57-58, 29-44 (2017). https://doi.org:10.1016/j.matbio.2016.10.003

11 Mao, M., Alavi, M. V., Labelle-Dumais, C. & Gould, D. B. Type IV Collagens and Basement Membrane Diseases: Cell Biology and Pathogenic Mechanisms. Curr Top Membr 76, 61–116 (2015). https://doi.org:10.1016/bs.ctm.2015.09.002

12 Kuo, D. S., Labelle-Dumais, C. & Gould, D. B. COL4A1 and COL4A2 mutations and disease: insights into pathogenic mechanisms and potential therapeutic targets. Hum Mol Genet 21, R97–R110 (2012). https://doi.org:10.1093/hmg/dds346

13 Mao, M. et al. Identification of fibronectin 1 as a candidate genetic modifier in a Col4a1 mutant mouse model of Gould syndrome. Dis Model Mech 14 (2021). https://doi.org:10.1242/dmm.048231

14 Iadecola, C. The Neurovascular Unit Coming of Age: A Journey through Neurovascular Coupling in Health and Disease. Neuron 96, 17–42 (2017). https://doi.org:10.1016/j.neuron.2017.07.030

15 Girouard, H. et al. Astrocytic endfoot Ca2+ and BK channels determine both arteriolar dilation and constriction. Proc Natl Acad Sci U S A 107, 3811–3816 (2010). https://doi.org:10.1073/pnas.0914722107

16 Petzold, G. C., Albeanu, D. F., Sato, T. F. & Murthy, V. N. Coupling of neural activity to blood flow in olfactory glomeruli is mediated by astrocytic pathways. Neuron 58, 897–910 (2008). https://doi.org:10.1016/j.neuron.2008.04.029

17 Iadecola, C. & Nedergaard, M. Glial regulation of the cerebral microvasculature. Nat Neurosci 10, 1369–1376 (2007). https://doi.org:10.1038/nn2003

18 Longden, T. A. et al. Capillary K(+)-sensing initiates retrograde hyperpolarization to increase local cerebral blood flow. Nat Neurosci 20, 717–726 (2017). https://doi.org:10.1038/nn.4533

19 Thakore, P. et al. Brain endothelial cell TRPA1 channels initiate neurovascular coupling. Elife 10 (2021). https://doi.org:10.7554/eLife.63040

20 Drew, P. J., Mateo, C., Turner, K. L., Yu, X. & Kleinfeld, D. Ultra-slow Oscillations in fMRI and Resting-State Connectivity: Neuronal and Vascular Contributions and Technical Confounds. Neuron 107, 782–804 (2020). https://doi.org:10.1016/j.neuron.2020.07.020

21 Kaplan, L., Chow, B. W. & Gu, C. H. Neuronal regulation of the blood-brain barrier and neurovascular coupling. Nat Rev Neurosci 21, 416–432 (2020). https://doi.org:10.1038/s41583-020-0322-2

22 Dabertrand, F. et al. PIP2 corrects cerebral blood flow deficits in small vessel disease by rescuing capillary Kir2.1 activity. Proc Natl Acad Sci U S A 118 (2021). https://doi.org:10.1073/pnas.2025998118

23 Huang, C. L., Feng, S. & Hilgemann, D. W. Direct activation of inward rectifier potassium channels by PIP2 and its stabilization by Gbetagamma. Nature 391, 803–806 (1998). https://doi.org:10.1038/35882

24 Hansen, S. B., Tao, X. & MacKinnon, R. Structural basis of PIP2 activation of the classical inward rectifier K+ channel Kir2.2. Nature 477, 495–U152 (2011). https://doi.org:10.1038/nature10370

25 Kuo, D. S. et al. Allelic heterogeneity contributes to variability in ocular dysgenesis, myopathy and brain malformations caused by Col4a1 and Col4a2 mutations. Hum Mol Genet 23, 1709–1722 (2014). https://doi.org:10.1093/hmg/ddt560

26 Cannistraro, R. J., et al. CNS small vessel disease: A clinical review. Neurology 92, 1146-1156 (2019). https://doi.org:10.1212/WNL.0000000000007654

27 Hilal, S. et al. Prevalence, risk factors and consequences of cerebral small vessel diseases: data from three Asian countries. J Neurol Neurosurg Psychiatry 88, 669–674 (2017). https://doi.org:10.1136/jnnp-2016-315324

28 Knot, H. J., Zimmermann, P. A. & Nelson, M. T. Extracellular K(+)-induced hyperpolarizations and dilatations of rat coronary and cerebral arteries involve inward rectifier K(+) channels. J Physiol 492 (Pt 2), 419–430 (1996). https://doi.org:10.1113/jphysiol.1996.sp021318

29 Dabertrand, F. et al. Potassium channelopathy-like defect underlies early-stage cerebrovascular dysfunction in a genetic model of small vessel disease. Proc Natl Acad Sci U S A 112, E796–805 (2015). https://doi.org:10.1073/pnas.1420765112

30 Wenceslau, C. F. et al. Guidelines for the measurement of vascular function and structure in isolated arteries and veins. American journal of physiology. Heart and circulatory physiology (2021). https://doi.org:10.1152/ajpheart.01021.2020

31 Jeanne, M., Jorgensen, J. & Gould, D. B. Molecular and Genetic Analyses of Collagen Type IV Mutant Mouse Models of Spontaneous Intracerebral Hemorrhage Identify Mechanisms for Stroke Prevention. Circulation 131, 1555–1565 (2015). https://doi.org:10.1161/CIRCULATIONAHA.114.013395

32 Yamasaki, E. et al. Faulty TRPM4 channels underlie age-dependent cerebral vascular dysfunction in Gould syndrome. Proc Natl Acad Sci U S A 120, e2217327120 (2023). https://doi.org:10.1073/pnas.2217327120

33 Girouard, H. et al. Astrocytic endfoot Ca2+ and BK channels determine both arteriolar dilation and constriction. P Natl Acad Sci USA 107, 3811–3816 (2010). https://doi.org:10.1073/pnas.0914722107

34 Park, L. et al. Age-dependent neurovascular dysfunction and damage in a mouse model of cerebral amyloid angiopathy. Stroke 45, 1815–1821 (2014). https://doi.org:10.1161/STROKEAHA.114.005179

35 Tarantini, S. et al. Pharmacologically-induced neurovascular uncoupling is associated with cognitive impairment in mice. J Cerebr Blood F Met 35, 1871–1881 (2015). https://doi.org:10.1038/jcbfm.2015.162

36 Kraeuter, A. K., Guest, P. C. & Sarnyai, Z. The Y-Maze for Assessment of Spatial Working and Reference Memory in Mice. Methods Mol Biol 1916, 105–111 (2019). https://doi.org:10.1007/978-1-4939-8994-2_10

37 Ohno, M. et al. BACE1 deficiency rescues memory deficits and cholinergic dysfunction in a mouse model of Alzheimer’s disease. Neuron 41, 27–33 (2004). https://doi.org:Doi10.1016/S0896-6273(03)00810-9

38 Oakley, H. et al. Intraneuronal beta-amyloid aggregates, neurodegeneration, and neuron loss in transgenic mice with five familial Alzheimer’s disease mutations: Potential factors in amyloid plaque formation. J Neurosci 26, 10129–10140 (2006). https://doi.org:10.1523/Jneurosci.1202-06.2006

39 Swonger, A. K. & Rech, R. H. Serotonergic and cholinergic involvement in habituation of activity and spontaneous alternation of rats in a Y maze. J Comp Physiol Psychol 81, 509–522 (1972). https://doi.org:10.1037/h0033690

40 Sarnyai, Z. et al. Impaired hippocampal-dependent learning and functional abnormalities in the hippocampus in mice lacking serotonin(1A) receptors. Proc Natl Acad Sci U S A 97, 14731–14736 (2000). https://doi.org:10.1073/pnas.97.26.14731

41 Mughal, A., Harraz, O. F., Gonzales, A. L., Hill-Eubanks, D. & Nelson, M. T. PIP2 Improves Cerebral Blood Flow in a Mouse Model of Alzheimer’s Disease. Function (Oxf) 2, zqab010 (2021). https://doi.org:10.1093/function/zqab010

42 Suer, S., Sickmann, A., Meyer, H. E., Herberg, F. W. & Heilmeyer, L. M., Jr. Human phosphatidylinositol 4-kinase isoform PI4K92. Expression of the recombinant enzyme and determination of multiple phosphorylation sites. Eur J Biochem 268, 2099–2106 (2001). https://doi.org:10.1046/j.1432-1327.2001.02089.x

43 Balla, A. & Balla, T. Phosphatidylinositol 4-kinases: old enzymes with emerging functions. Trends Cell Biol 16, 351–361 (2006). https://doi.org:10.1016/j.tcb.2006.05.003

44 Gehrmann, T. et al. Functional expression and characterisation of a new human phosphatidylinositol 4-kinase PI4K230. Biochim Biophys Acta 1437, 341–356 (1999). https://doi.org:10.1016/s1388-1981(99)00029-3

45 Harraz, O. F., Longden, T. A., Dabertrand, F., Hill-Eubanks, D. & Nelson, M. T. Endothelial GqPCR activity controls capillary electrical signaling and brain blood flow through PIP2 depletion. P Natl Acad Sci USA 115, E3569–E3577 (2018). https://doi.org:10.1073/pnas.1800201115

46 Mishra, R., Patel, H., Alanazi, S., Kilroy, M. K. & Garrett, J. T. PI3K Inhibitors in Cancer: Clinical Implications and Adverse Effects. Int J Mol Sci 22 (2021). https://doi.org:10.3390/ijms22073464

47 Zagaglia, S. et al. Neurologic phenotypes associated with COL4A1/2 mutations: Expanding the spectrum of disease. Neurology 91, e2078–e2088 (2018). https://doi.org:10.1212/WNL.0000000000006567

48 Meuwissen, M. E. et al. The expanding phenotype of COL4A1 and COL4A2 mutations: clinical data on 13 newly identified families and a review of the literature. Genet Med 17, 843–853 (2015). https://doi.org:10.1038/gim.2014.210

49 Jeanne, M. & Gould, D. B. Genotype-phenotype correlations in pathology caused by collagen type IV alpha 1 and 2 mutations. Matrix Biol 57-58, 29-44 (2017). https://doi.org:10.1016/j.matbio.2016.10.003

50 Kuo, D. S., Labelle-Dumais, C. & Gould, D. B. COL4A1 and COL4A2 mutations and disease: insights into pathogenic mechanisms and potential therapeutic targets. Hum Mol Genet 21, R97–110 (2012). https://doi.org:10.1093/hmg/dds346

51 Labelle-Dumais, C. et al. COL4A1 Mutations Cause Neuromuscular Disease with Tissue-Specific Mechanistic Heterogeneity. Am J Hum Genet 104, 847–860 (2019). https://doi.org:10.1016/j.ajhg.2019.03.007

52 Verzijl, N. et al. Effect of collagen turnover on the accumulation of advanced glycation end products. J Biol Chem 275, 39027–39031 (2000). https://doi.org:10.1074/jbc.M006700200

53 Nissen, R., Cardinale, G. J. & Udenfriend, S. Increased turnover of arterial collagen in hypertensive rats. Proc Natl Acad Sci U S A 75, 451–453 (1978). https://doi.org:10.1073/pnas.75.1.451

54 Mays, P. K., McAnulty, R. J., Campa, J. S. & Laurent, G. J. Age-related changes in collagen synthesis and degradation in rat tissues. Importance of degradation of newly synthesized collagen in regulating collagen production. Biochem J 276 (Pt 2), 307–313 (1991). https://doi.org:10.1042/bj2760307

55 Boot-Handford, R. P., Kurkinen, M. & Prockop, D. J. Steady-state levels of mRNAs coding for the type IV collagen and laminin polypeptide chains of basement membranes exhibit marked tissue-specific stoichiometric variations in the rat. J Biol Chem 262, 12475–12478 (1987).

56 Mao, M., Labelle-Dumais, C., Tufa, S. F., Keene, D. R. & Gould, D. B. Elevated TGFbeta signaling contributes to ocular anterior segment dysgenesis in Col4a1 mutant mice. Matrix Biol 110, 151–173 (2022). https://doi.org:10.1016/j.matbio.2022.05.001

57 Branyan, K. et al. Elevated TGFbeta Signaling Contributes to Cerebral Small Vessel Disease in Mouse Models of Gould Syndrome. Matrix Biol (2022). https://doi.org:10.1016/j.matbio.2022.11.007

58 Bakin, A. V., Tomlinson, A. K., Bhowmick, N. A., Moses, H. L. & Arteaga, C. L. Phosphatidylinositol 3-kinase function is required for transforming growth factor beta-mediated epithelial to mesenchymal transition and cell migration. Journal of Biological Chemistry 275, 36803–36810 (2000). https://doi.org:DOI10.1074/jbc.M005912200

59 Shin, I., Bakin, A. V., Rodeck, U., Brunet, A. & Arteaga, C. L. Transforming growth factor beta enhances epithelial cell survival via Akt-dependent regulation of FKHRL1. Mol Biol Cell 12, 3328–3339 (2001). https://doi.org:10.1091/mbc.12.11.3328

60 Hara, K. et al. Association of HTRA1 mutations and familial ischemic cerebral small-vessel disease. N Engl J Med 360, 1729–1739 (2009). https://doi.org:10.1056/NEJMoa0801560

61 Favor, J. et al. Type IV procollagen missense mutations associated with defects of the eye, vascular stability, the brain, kidney function and embryonic or postnatal viability in the mouse, Mus musculus: an extension of the Col4a1 allelic series and the identification of the first two Col4a2 mutant alleles. Genetics 175, 725-736 (2007). https://doi.org:10.1534/genetics.106.064733

