## Supplementary figures and images for "PI3K block restores age-dependent neurovascular coupling defects associated with cerebral small vessel disease"

### Fig. 1 - supplement 1

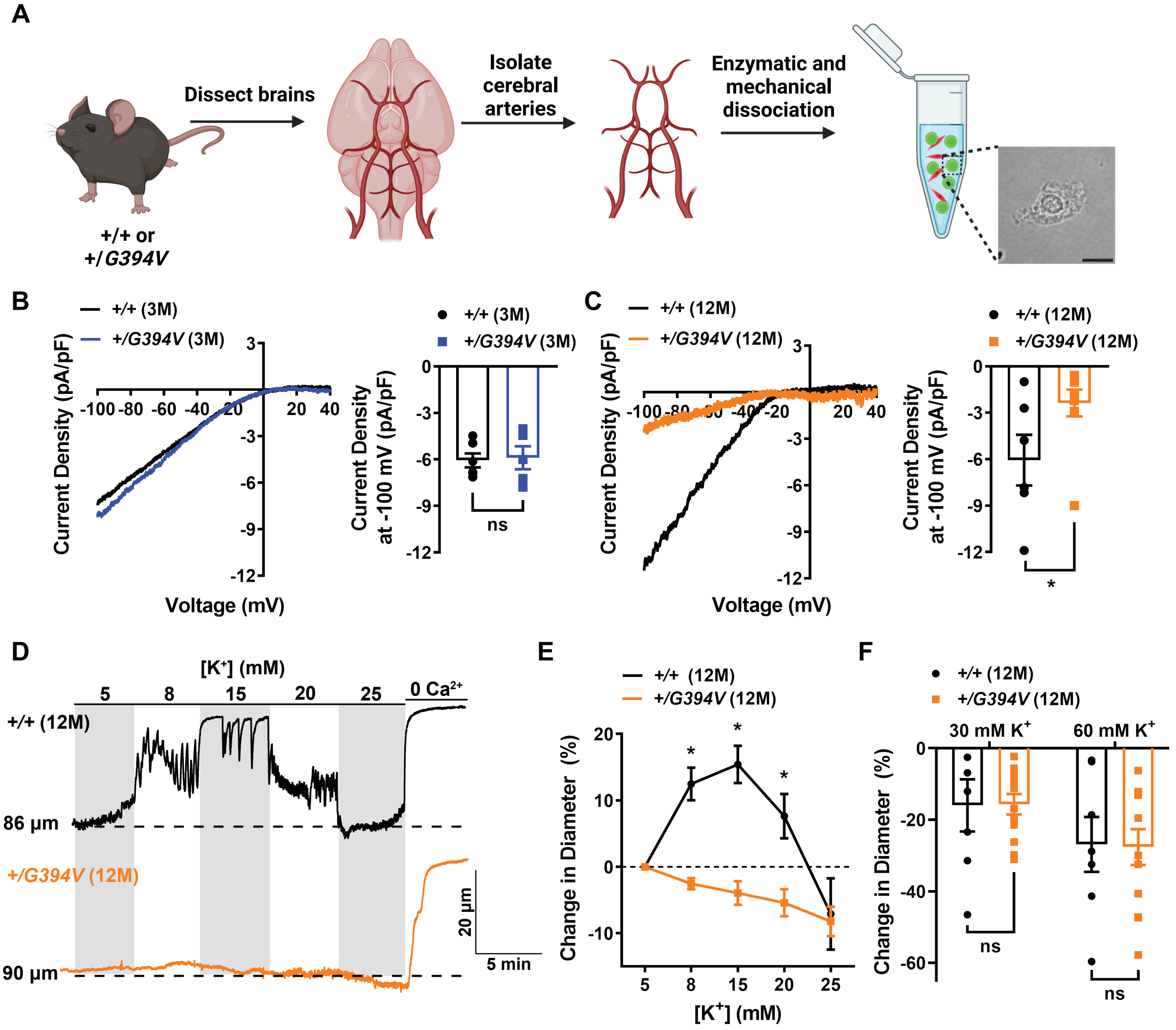

### Fig. 1 - supplement 2

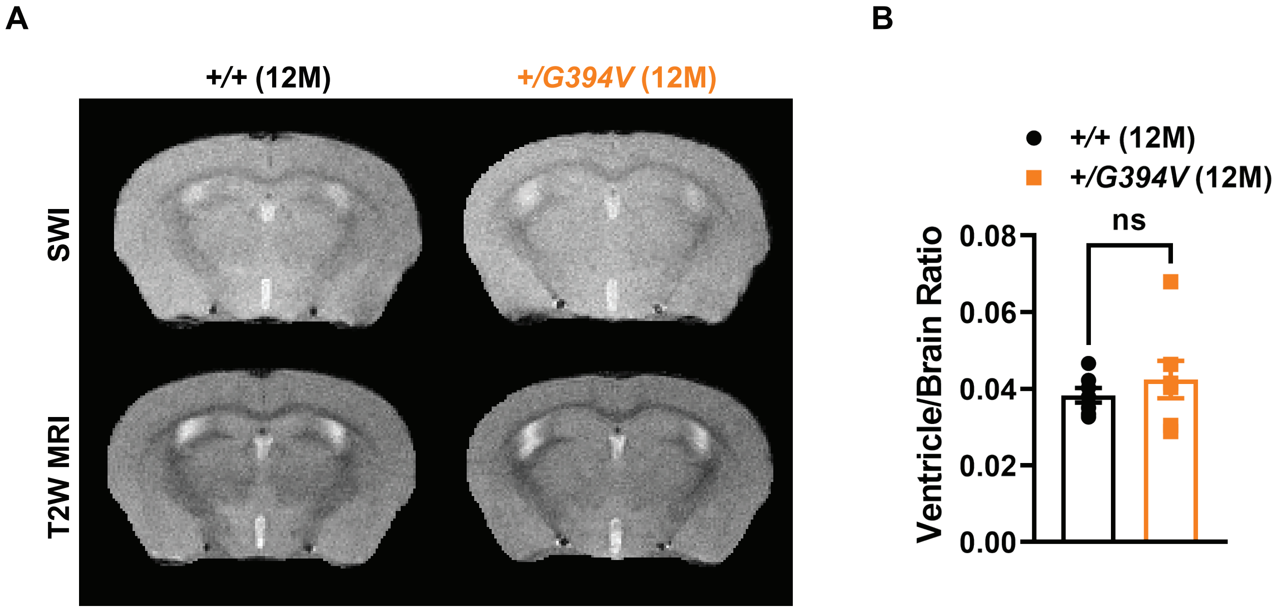

### Fig. 2 - supplement 1

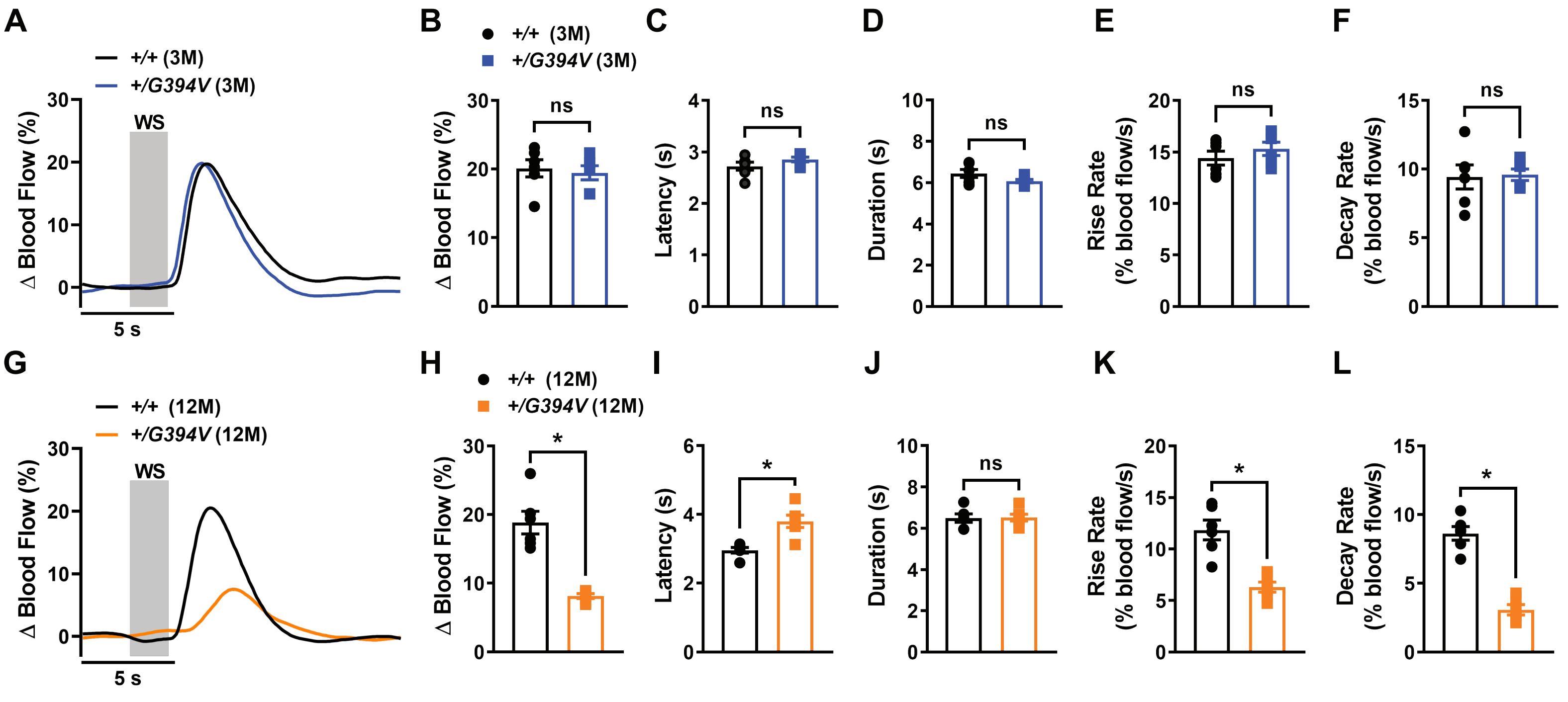

### Fig. 2 - supplement 2

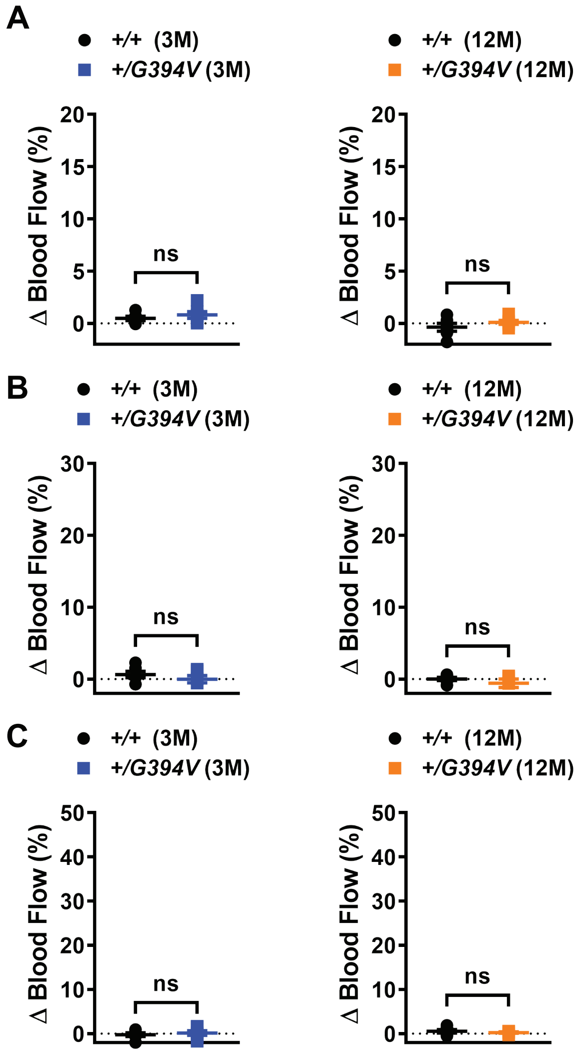

### Fig. 4 - supplement 1

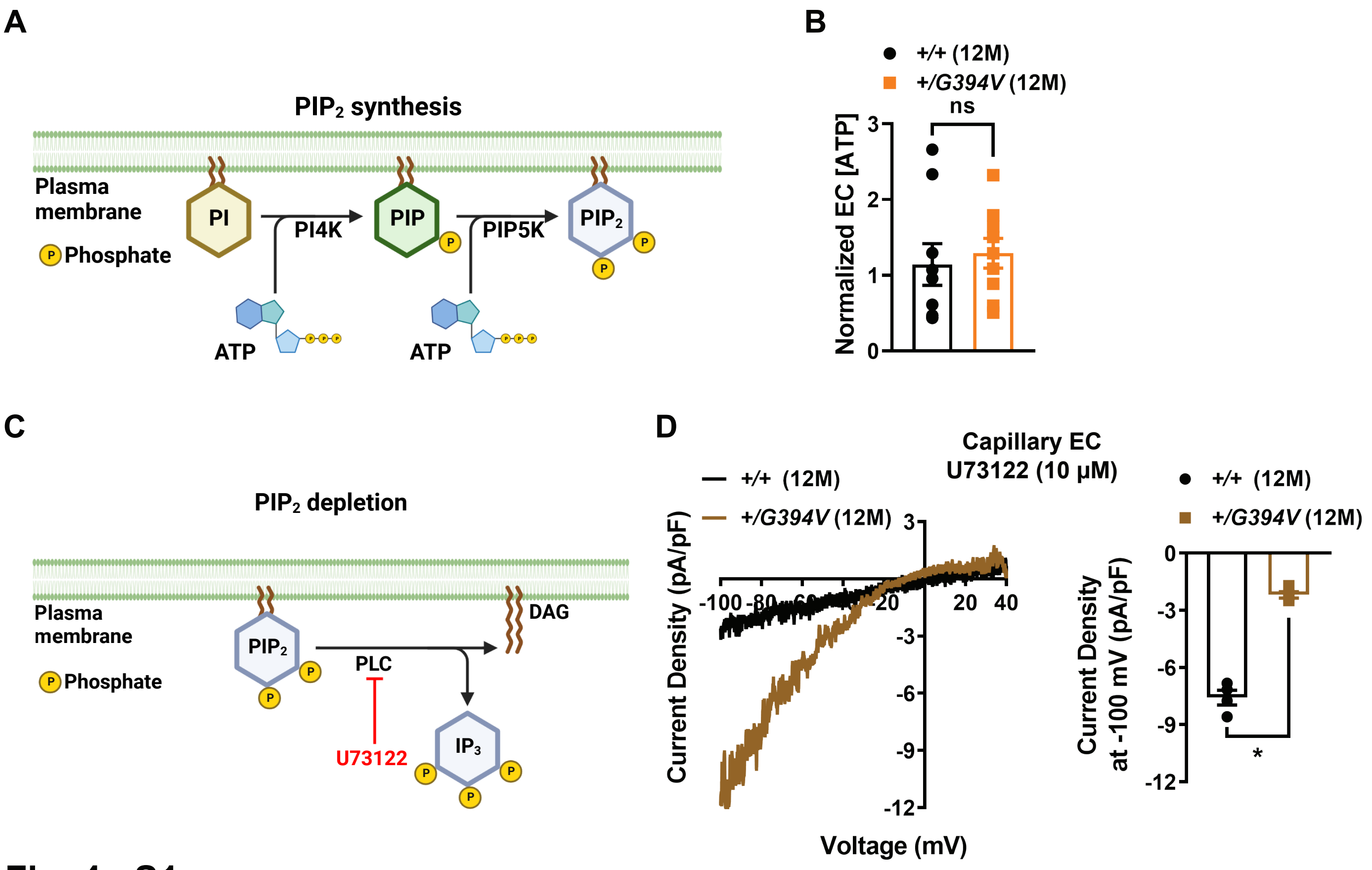

### Fig. 4 - supplement 2

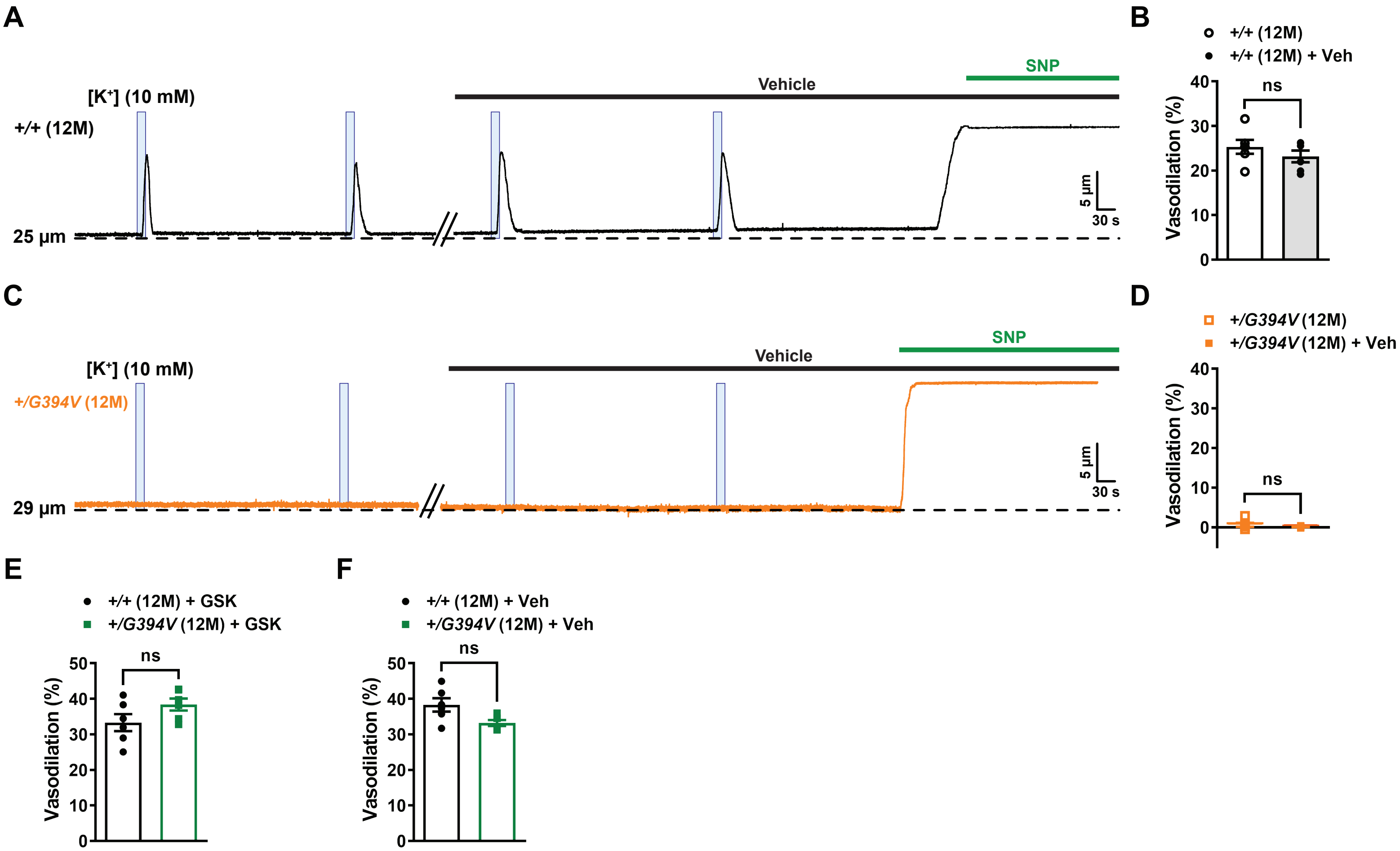

### Fig. 5 - supplement 1

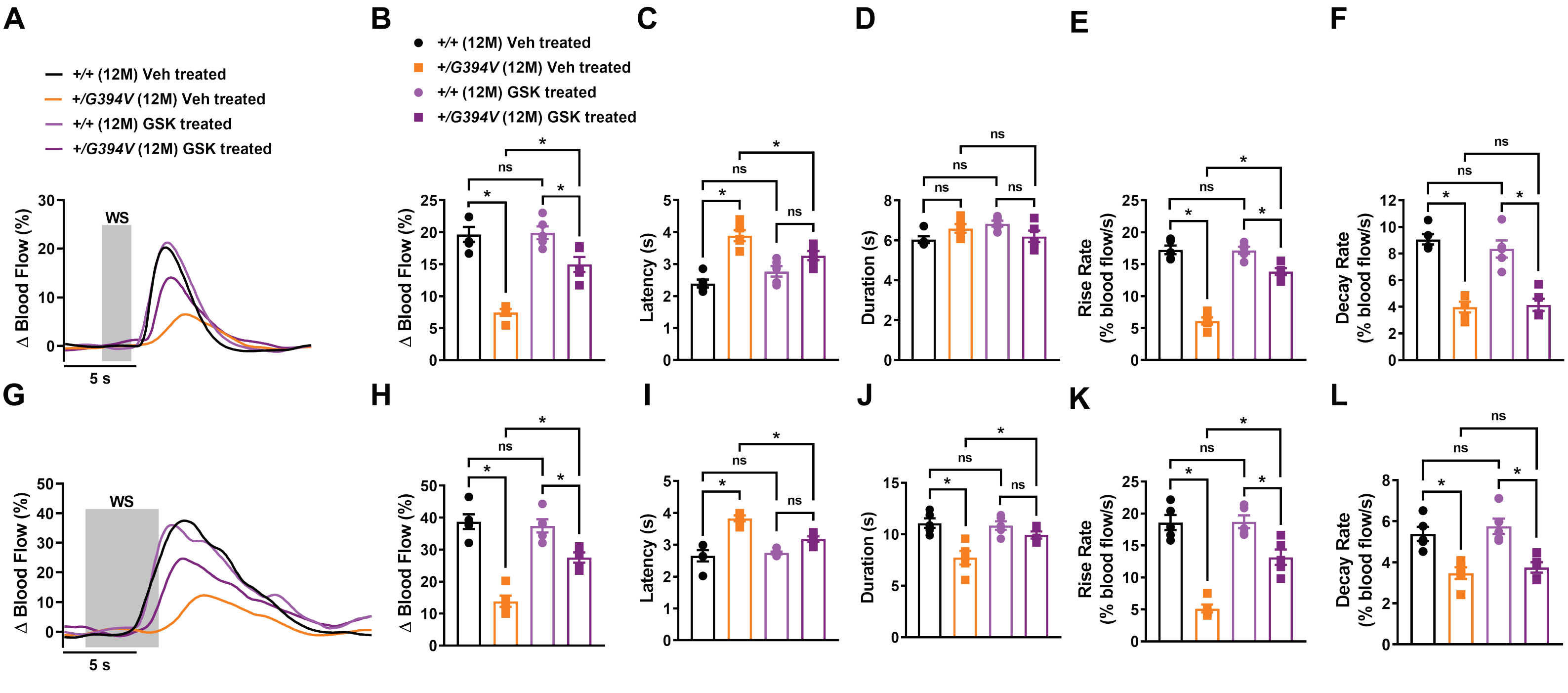

### Fig. 5 - supplement 2

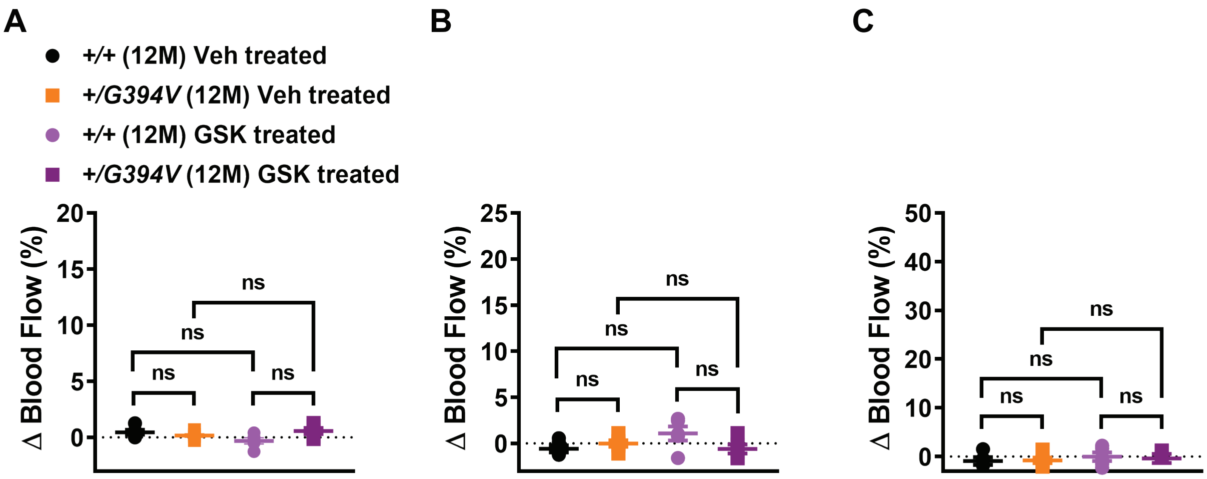
